# Modality-chain reasoning enables multimodal protein modelling and design

**DOI:** 10.1101/2025.07.21.665832

**Authors:** Chaozhong Liu, Linlin Chao, Shaomin Ji, Hao Wang, Guowei Zhou, Jiayou Zheng, Dubeiqi Hong, Yucheng Guo, Taorui Jiang, Zhangyang Gao, Ming Yang, Jianfeng Pan, Stan Z. Li, Xiaoming Zhang

**Author notes:** Equal Contribution (Chaozhong Liu, Linlin Chao). Work done during internship at BioMap (Jiayou Zheng, Taorui Jiang).

## Abstract

Reasoning has emerged as a central capability of large language models, yet how it should be formulated for scientific foundation models remains unclear because scientific knowledge is distributed across interdependent, domain-specific representations. Here we introduce modality-chain reasoning, which organizes representations into ordered computational chains, each conditioning prediction or generation of the next. Based on this principle, we develop ProteinReasoner, a multimodal generative protein foundation model that sequentially connects amino acid sequence, evolutionary constraints and three-dimensional structure within a shared autoregressive architecture. Across zero-shot structure prediction, inverse folding and fitness prediction, ProteinReasoner outperformed two multimodal protein foundation models, while controlled comparisons supported the functional contribution of the modality chain. We further extended this principle beyond pretraining: reorganizing the chain across successive structural states enabled multiple-conformation prediction, while introducing experimental feedback as an additional modality enabled an in-context learning paradigm for protein optimization without target-specific parameter updates. In particular, across thermostability and affinity-maturation evaluations, this paradigm improved over matched fine-tuned models and showed stronger mean performance than target-specific active-learning baselines. These results establish modality-chain reasoning as a unified and effective foundation-modelling strategy in protein science. More broadly, they suggest a general route towards reasoning across interdependent representations in other scientific domains.

## 1. Introduction

Reasoning has emerged as a central capability of large language models, substantially improving their performance on complex tasks^1,2^. An analogous capability could be valuable for scientific foundation models by enabling them to integrate complementary knowledge across tasks and support both prediction and generation. However, scientific foundation models differ from language models in that scientific knowledge is represented not only in text, but also through multiple domain-specific representations of the same system. These representations are often interdependent, such that information in one can constrain the plausible states of another. How reasoning should be formulated across these heterogeneous scientific representations remains unclear.

Proteins provide a natural setting in which to examine this question. A protein can be described through its amino acid sequence, evolutionary variation, three-dimensional structure and diverse structural, biophysical and functional properties. These representations capture complementary aspects of the same molecular system and are linked by physical, evolutionary and functional constraints. In structure prediction, as exemplified by AlphaFold2^3^, MSAs summarize residue conservation and covariation, providing evolutionary constraints that guide the inference of three-dimensional structure^4,5^ (**Fig. 1a**). Protein optimization^6^ follows a related progression: experimentally measured mutant sequences collectively provide a representation of sequence– function relationships that guides the design of subsequent mutant sequences (**Fig. 1b**). These workflows suggest that reasoning in protein science can be viewed as progression across interdependent representations, with each stage constraining the plausible outcomes of the next.

**Fig. 1.**
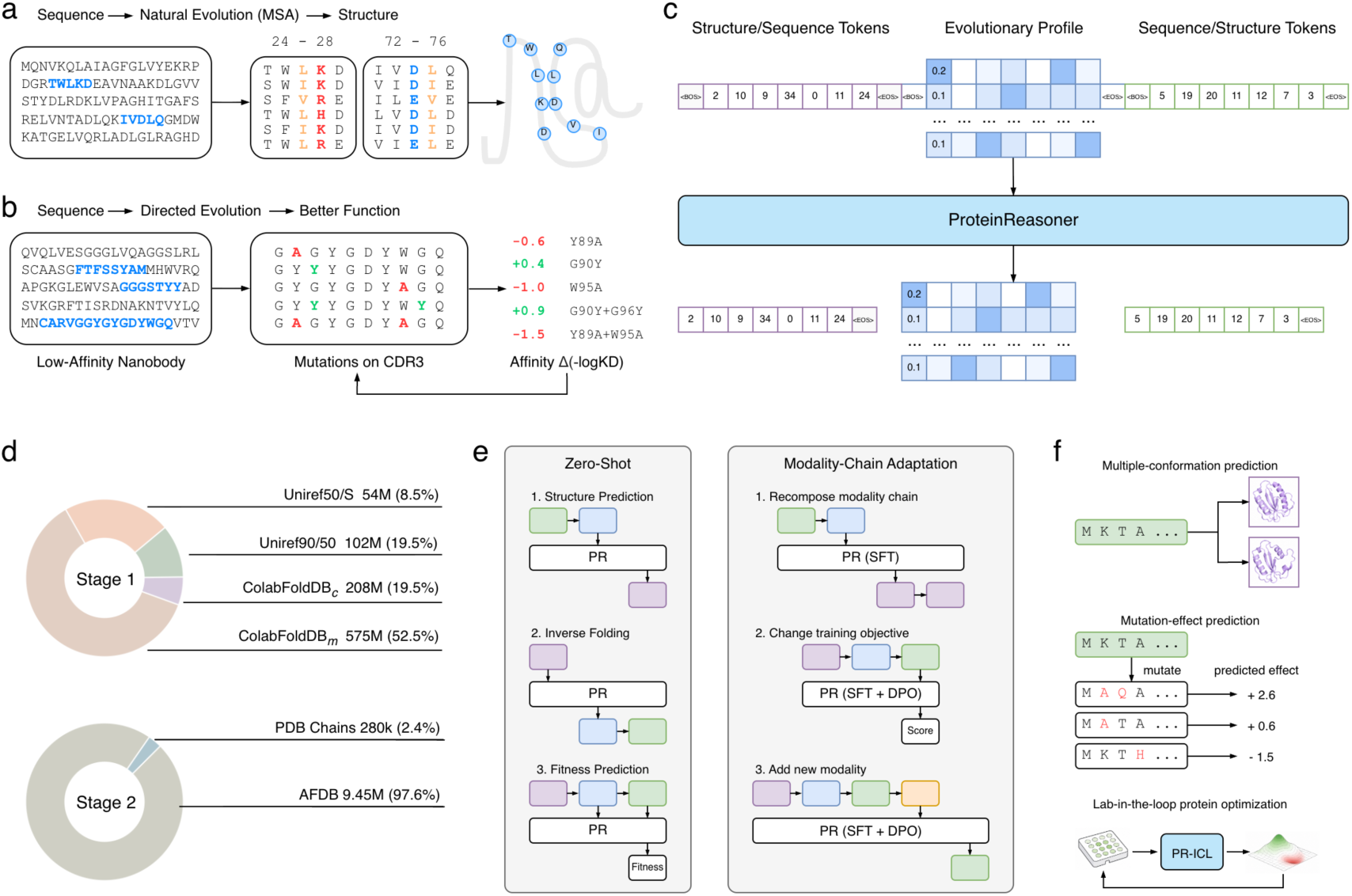
ProteinReasoner implements modality-chain reasoning. **a,** Natural evolutionary information encoded by multiple sequence alignments (MSAs) constrains structure inference from sequence. **b**, Wet-lab measurements of mutant fitness guide the design of subsequent mutant sequences during protein optimization. **c**, ProteinReasoner architecture and multimodal pretraining. Sequence tokens, evolutionary profiles and structure tokens are represented as distinct modalities and processed by a shared autoregressive transformer. ‘Structure/Sequence Tokens’ and ‘Sequence/Structure Tokens’ denote the two chain directions. **d**, Statistics of the two-stage pretraining datasets. Stage 1 uses sequence-only datasets, whereas Stage 2 uses proteins with associated structural and evolutionary information. Percentages indicate contributions to the total number of training tokens. **e**, Zero-shot applications and general strategies for task adaptation, including recomposition of modality chains, modification of training objectives and addition of new modalities. **f**, Downstream applications evaluated in this study: multiple-conformation prediction, mutation-effect prediction and lab-in-the-loop protein optimization with in-context learning.

Protein language models provide a natural computational foundation for learning such relationships at scale. Sequence-based models, including the ESM family^7,8^, have learned representations that support diverse tasks such as structure prediction, mutation-effect estimation and function annotation^7–9^. Their success has motivated multimodal protein foundation models, including ESM3^10^, ProSST^11^, SaProt^12^, and DPLM-2^13^, which incorporate structural or evolutionary information alongside sequence. However, these models generally combine modalities through joint representations or direct mappings between inputs and outputs. Such formulations can exploit complementary information, but leave the ordered dependencies that organize protein-science workflows largely implicit. We therefore asked whether explicitly organizing protein representations into ordered reasoning chains could provide a unified and effective foundation-modelling strategy across diverse protein prediction and generation tasks.

To address this question, we introduce modality-chain reasoning, a framework that organizes modality-specific representations into ordered computational chains, such that each representation conditions the prediction or generation of the next. We instantiate this principle in ProteinReasoner, a multimodal generative protein foundation model that connects sequence, MSA-derived evolutionary constraints and structure within a shared autoregressive architecture (**Fig. 1c**). We first evaluated whether modality-chain reasoning supports effective multimodal protein modelling through zero-shot structure prediction, inverse folding and fitness prediction, together with controlled comparisons against direct end-to-end modelling. We then tested its extensibility through downstream reformulations: reorganizing the chain across successive structural states for multiple-conformation prediction, and introducing experimental feedback as an additional modality for iterative protein optimization (**Fig. 1e,f**). The experimental-feedback extension establishes an in-context learning paradigm in which a shared foundation model can adapt to target-specific sequence–property relationships without target-specific parameter updates.

## 2. Results

### 2.1 ProteinReasoner implements modality-chain reasoning across protein modalities

ProteinReasoner is a multimodal generative protein foundation model that implements modality-chain reasoning across amino acid sequence, evolutionary constraints and three-dimensional structure (**Fig. 1c**; **Extended Data Fig. 1a**). Here, modality-chain reasoning refers to keeping scientific representations explicit, arranging them according to task-relevant dependencies, and modelling them sequentially so that each representation conditions the prediction or generation of the next. Accordingly, ProteinReasoner preserves these information sources as distinct modality segments within a shared autoregressive context rather than combining them into a single input representation.

The input chain to ProteinReasoner consists of three modalities: amino acid sequence, evolutionary profile and three-dimensional structure. The amino acid sequence is tokenized at residue resolution as a length-*L* sequence, whereas structure is discretized into residue-aligned tokens using the DPLM-2 structure tokenizer^13^. Evolutionary constraints are represented as an evolutionary profile (**Extended Data Fig. 1b**), an *L* × 21 matrix derived from MSAs in which each row gives the frequencies of the 20 standard amino acids and a gap at one residue position, summarizing position-specific evolutionary preferences.

During pretraining, the three modalities are arranged in two directional chains: sequence → evolutionary profile→ structure and structure → evolutionary profile → sequence (**Extended Data Fig. 1a**, bottom). ProteinReasoner implements these chains using a shared decoder-only Transformer backbone with modality-specific input projections, output heads and boundary tokens. The output heads predict amino acid tokens, evolutionary profile vectors and structure tokens position by position. These chains correspond to folding and inverse folding, respectively, training the model to propagate information through the evolutionary profile in either direction. The profile thereby serves as an explicit intermediate representation linking sequence and structure.

ProteinReasoner was pretrained using a two-stage strategy (**Fig. 1d**). In the first stage, a sequence-only language model^14^ was trained on approximately one trillion amino acid tokens. In the second stage, this model was further trained on a multimodal dataset comprising 9.45 million protein structures from the AlphaFold Database^15^ and 279,921 experimentally resolved chains from PDB-REDO^16^ with refined X-ray structures from the Protein Data Bank^17^. We trained models with 150 million, 650 million, and 3 billion parameters, using up to 1.89×10^11^ multimodal tokens. On held-out validation data, larger models achieved lower sequence and structure perplexity and lower Kullback–Leibler divergence for predicted evolutionary profiles (**Extended Data Fig. 1c**).

### 2.2 Modality-chain reasoning supports effective zero-shot protein modelling

To assess whether modality-chain reasoning provides an effective paradigm for protein modelling, we evaluated ProteinReasoner under zero-shot settings on three representative tasks: structure prediction^18^, inverse folding^19^, and fitness prediction^20^. We selected ESM3-Open-1.4B and DPLM-2 as complementary baselines: ESM3 provides a general-purpose multimodal protein foundation model with broad generative capabilities, whereas DPLM-2 is particularly relevant because ProteinReasoner adopts its structure tokenizer. All models were evaluated using the benchmark datasets and protocols described in Methods.

For structure prediction and inverse folding, we first evaluated test proteins absent from the ProteinReasoner pretraining corpus. This exact-overlap control provides a practical basis for comparison because the broad pretraining sources of ESM3 and DPLM-2, including AlphaFoldDB^15^ and ESM Atlas^8^ for ESM3 and PDB-derived data for DPLM-2, make the exposure of individual benchmark proteins difficult to determine. The evaluation included the combined CASP14, CASP15 and CAMEO set (n = 204) and the PDB holdout set (n = 449). To further assess generalization under stringent novelty constraints, we pooled these benchmarks and retained proteins sharing less than 50% (n = 199) or 30% (n = 88) sequence identity with any ProteinReasoner pretraining sequence. These similarity restrictions were applied only to ProteinReasoner because the complete training sequences of the baseline models were unavailable.

In structure prediction, ProteinReasoner was evaluated in two configurations: an externally guided setting, in which an MSA-derived evolutionary profile was provided, and an internally inferred setting, in which the model generated the profile before predicting structure (**Fig. 2a**). Predicted structure tokens were decoded into atomic coordinates and evaluated using root-mean-square deviation (RMSD) and Template Modeling score (TM-score)^21^. In the externally guided setting, ProteinReasoner consistently achieved the strongest mean performance across the full and similarity-controlled benchmarks (Fig. 2b,c), reducing RMSD by 5.3–19.2% and increasing TM-score by 0.9–7.9% relative to the strongest baseline. Pairwise comparisons supported these gains for both metrics on the CAMEO/CASP, PDB holdout and <50% sequence-identity benchmarks; on the <30% subset, RMSD remained significantly lower whereas the TM-score difference was not significant. Performance generally improved with model scale in both settings. Although externally provided profiles remained more effective than internally generated ones, ProteinReasoner-3B approached ESM3 without external MSA input: differences were not significant in six of eight benchmark–metric comparisons, with lower RMSD on the PDB holdout but lower TM-score on CAMEO/CASP. These results support the evolutionary profile as an effective intermediate representation for sequence-to-structure modelling, while indicating that the utility of internally generated profiles increases with model capacity.

**Fig. 2.**
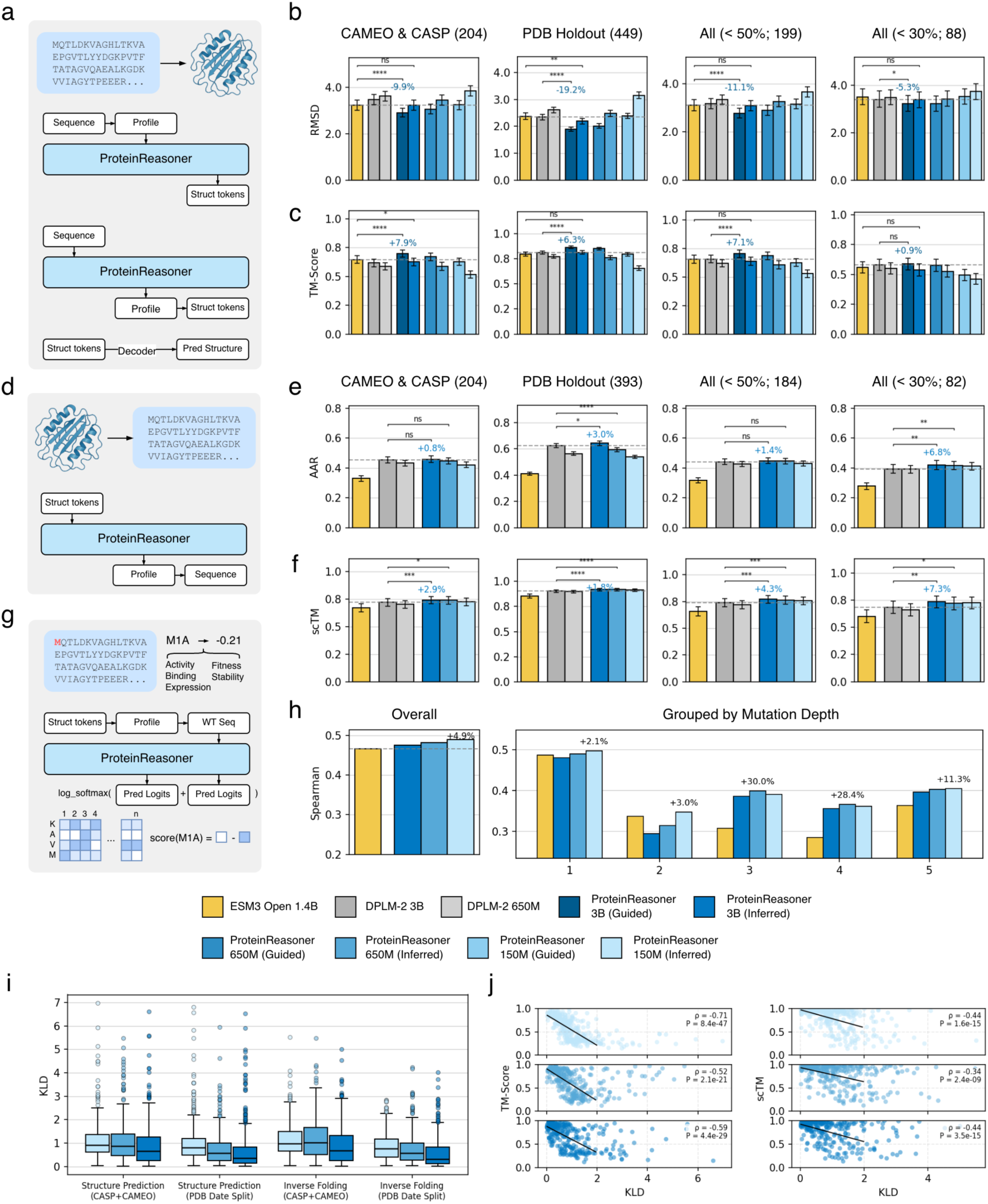
Zero-shot evaluation and analysis of modality-chain reasoning in ProteinReasoner. **a**, Structure prediction in externally guided and internally inferred settings. In the guided setting, an MSA-derived evolutionary profile is provided; in the inferred setting, ProteinReasoner first generates the profile before predicting structure tokens, which are decoded using the DPLM-2 structure decoder. **b,c**, Structure prediction performance measured by RMSD (**b**) and TM-score (**c**) on the combined CAMEO and CASP set, the PDB holdout set, and pooled subsets with <50% or <30% sequence identity to the pretraining data. **d**, Inverse protein folding in the internally inferred setting, in which ProteinReasoner generates an evolutionary profile and amino acid sequence from structure tokens. **e,f**, Inverse folding performance measured by amino acid recovery (AAR; **e**) and self-consistency TM-score (scTM; **f**) on the corresponding benchmark sets. **g**, Fitness prediction with the inverse-folding chain as input. ProteinReasoner combines log-softmax-normalized sequence- and profile-head logits and scores each mutation by the log-odds difference between the mutant and WT amino acids. **h**, ProteinGym fitness prediction performance, shown as the overall Spearman correlation (left) and stratified by mutation depth (right). **i**, Quality of internally inferred evolutionary profiles, measured by Kullback–Leibler divergence (KLD), across structure prediction and inverse folding tasks. **j**, Relationship between profile KLD and downstream performance for structure prediction, measured by TM-score (left), and inverse folding, measured by scTM (right). Each point represents one protein, and lines show linear fits using proteins with KLD ≤ 2. Spearman’s ρ and two-sided *P* values are shown in each panel. Percentages above bars indicate the relative performance change of the best-performing ProteinReasoner configuration compared with the best-performing baseline. In **b,c,e,f**, bars show mean performance and error bars indicate 95% bootstrap confidence intervals across proteins. Statistical brackets in **b,c** compare the best-performing ProteinReasoner and external baseline configurations and ProteinReasoner-3B inferred with ESM3-Open-1.4B; those in **e,f** compare the best-performing ProteinReasoner and external baseline configurations and ProteinReasoner-650M with DPLM-2-3B. Statistical significance was assessed using two-sided paired Wilcoxon signed-rank tests. *P* values are unadjusted. ns, *P* ≥ 0.05; *, *P* < 0.05; **, *P* < 0.01; ***, *P* < 0.001; ****, *P* < 0.0001.

We next evaluated the reverse modality chain through inverse folding, where the model generates sequences compatible with a given structure. ProteinReasoner was assessed only in the internally inferred setting to avoid leaking sequence information through an MSA-derived profile (**Fig. 2d**). Performance was measured by amino acid recovery (AAR) and self-consistency TM-score (scTM), which quantify native-sequence recovery and the structural consistency between the target structure and structures predicted by AlphaFold2^3^ from the generated sequences, respectively. Across the full and similarity-controlled benchmarks, the best-performing ProteinReasoner configurations achieved higher mean AAR and scTM than the ESM3 and DPLM-2 baselines (**Fig. 2e,f**), with improvements of up to 6.8% and 7.3%, respectively, relative to the strongest baseline. The scTM advantage was significant across all four benchmark sets, whereas the AAR advantage was significant on the PDB holdout and <30% sequence-identity subset. Notably, ProteinReasoner-650M achieved significantly higher scTM than DPLM-2-3B across all four benchmarks despite its smaller model size, whereas AAR differences between these models were benchmark-dependent. Scaling from 150M to 650M parameters generally improved performance, whereas further gains at 3B were modest. Thus, the reverse modality chain supports generalizable zero-shot sequence generation with strong compatibility to the target structure.

Finally, we assessed whether representations learned through modality-chain pretraining support mutation-effect prediction using the ProteinGym DMS substitution dataset^22^ (n = 217). Inputs followed the inverse-folding order, comprising structure, evolutionary profile and wild-type sequence (**Fig. 2g**), and fitness scores were derived by combining predictions from the profile and sequence output heads (see **Methods**). Across the ProteinGym benchmark, ProteinReasoner improved Spearman correlation by 4.9% relative to ESM3-Open-1.4B (**Fig. 2h**). Performance did not improve monotonically with model scale, with the strongest overall result obtained by the 150M model. This is consistent with recent evidence that model scale and performance are not strictly proportional across ProteinGym benchmarks^23^. When benchmarks were grouped by mutation depth, ProteinReasoner matched or outperformed ESM3 across all groups, with differences of up to 30.0% at intermediate-to-high mutation depths. These results extend the zero-shot utility of modality-chain pretraining beyond generative tasks to mutation-effect prediction across increasing mutational complexity.

These evaluations show that the same modality-chain-pretrained framework achieves strong zero-shot performance in both generative and predictive protein tasks and in both chain directions. Benchmark performance alone, however, does not establish whether organizing complementary modalities into an ordered chain itself contributes to these gains.

To test this directly, we compared ProteinReasoner with an end-to-end ablation trained directly between sequence and structure and a uniform-profile ablation in which the evolutionary profile was replaced by a non-informative matrix (**Extended Data Fig. 2b, c**). In structure prediction, the internally inferred ProteinReasoner outperformed the end-to-end model when both received only the amino acid sequence as input, with significantly higher TM-score across all four benchmarks and significantly lower RMSD on the PDB holdout and both sequence-identity-controlled subsets (**Extended Data Fig. 2d,e**). Replacing the evolutionary profile with a uniform matrix significantly reduced performance for both metrics across all four benchmarks, showing that the benefit depends on task-relevant evolutionary information rather than simply adding another segment to the chain.

The benefit of the complete sequence–profile–structure formulation was also observed in fitness prediction (**Extended Data Fig. 2h**). In inverse folding, ProteinReasoner achieved significantly lower AAR than the end-to-end model on three of the four benchmarks, indicating greater sequence diversity. scTM remained similar on three benchmarks, with only a small decrease on the PDB holdout reaching significance (**Extended Data Fig. 2f,g**), indicating that this increased diversity was accompanied by largely preserved compatibility with the target structure. Together, these ablations support modality-chain reasoning as a modelling strategy in which biologically meaningful intermediate representations provide task-relevant conditioning for subsequent modelling. Complete results and additional ablations of pretraining design choices are provided in the **Supplementary Information**.

To further examine the contribution of the intermediate evolutionary modality, we analysed model-generated profile quality and its relationship with downstream performance. Profile quality, measured by Kullback– Leibler divergence (KLD) from reference MSA-derived profiles, improved with model scale (**Fig. 2i**). Across both structure prediction and inverse folding, profile KLD was significantly negatively correlated with downstream performance (**Fig. 2j**), further linking the quality of the inferred intermediate representation to subsequent modelling performance.

We next assessed how the model incorporates the evolutionary profile during inference. When attention weights were averaged across all layers and heads, approximately 17% of cross-modality attention from the subsequent modality was directed to the profile in both structure prediction and inverse folding (**Extended Data Fig. 3a**). We then examined typical attention heads with high target-to-profile attention. In structure prediction, these heads assigned greater attention to profiles derived from deeper MSAs, whereas in inverse folding, attention increased as the KLD of the generated profile decreased (**Extended Data Fig. 3b**). Although attention weights do not establish causality, these patterns are consistent with the model conditioning subsequent generation on the intermediate profile and varying its reliance with profile quality. These analyses provide complementary evidence that downstream modelling is sensitive to both the quality and task-dependent use of the intermediate representation.

### 2.3 Extending modality-chain reasoning to multiple-conformation prediction

To assess whether modality-chain reasoning can serve as a general strategy for protein modelling beyond the pretraining tasks, we applied ProteinReasoner to multiple-conformation prediction^24^, which aims to recover distinct experimentally observed structural states of the same protein from a shared sequence (**Fig. 3a**). Predicting alternative conformations is important for understanding protein function, as conformational changes mediate functional states, molecular interactions and regulation. Existing approaches predominantly rely on sampling strategies, including diffusion- and flow-based generative models^25–27^, and AlphaFold-based methods that induce diversity through MSA perturbations or inference settings^28–30^.

**Fig. 3.**
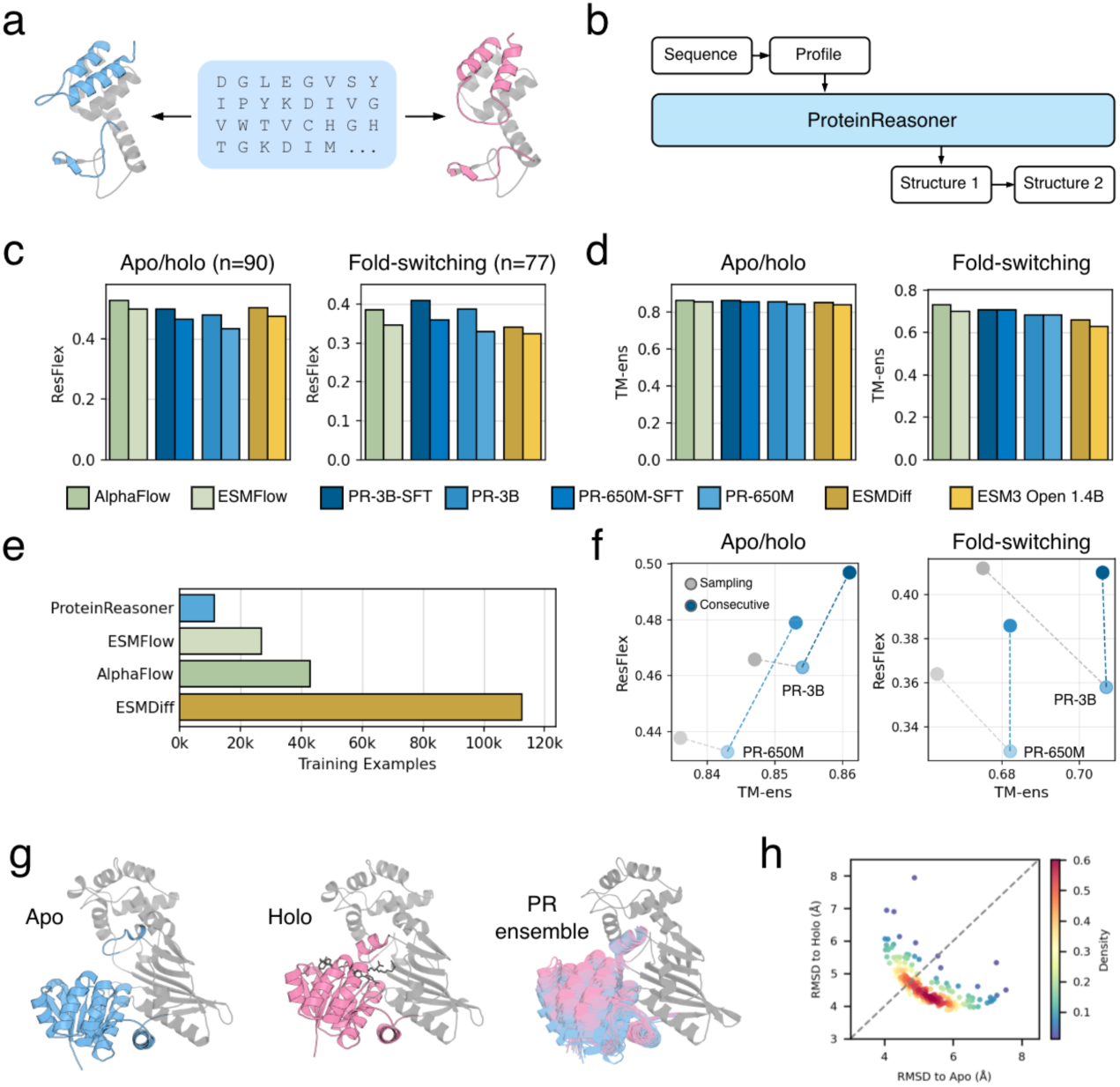
ProteinReasoner extends modality-chain reasoning to multiple-conformation prediction. **a**, Schematic of multiple-conformation prediction, in which a single protein sequence can adopt distinct structural states. **b**, Extended modality chain for multiple-conformation prediction. Given a sequence and evolutionary profile, ProteinReasoner generates two structures sequentially, allowing the second structure to condition on the first. **c,d**, Performance on the apo/holo (left) and fold-switching (right) benchmarks, evaluated by residue flexibility (ResFlex; **c**) and ensemble TM-score (TM-ens; **d**). **e**, Number of training conformations used by the benchmarked models. **f**, Comparison of consecutive generation with the three-modality sampling strategy on the apo/holo (left) and fold-switching (right) benchmarks. Dashed lines connect pretrained and fine-tuned models. Points show mean performance. **g**, Case study of saccharopine reductase from *Magnaporthe grisea*. The apo structure (PDB 1E5L), holo structure (PDB 1E5Q) and a 20-structure ensemble generated by fine-tuned ProteinReasoner-3B are shown from left to right. Domain III and the flexible loop are coloured blue in the apo state and pink in the holo state, with the remaining structure shown in grey. Ensemble structures are coloured according to their similarity to the apo and holo states. **h**, Distribution of 200 ProteinReasoner-predicted conformations for saccharopine reductase. Each point represents one predicted structure, plotted by its Cα RMSD to the apo structure on the x axis and the holo structure on the y axis; color indicates local density. PR, ProteinReasoner; SFT, supervised fine-tuning.

Rather than treating multiple-conformation prediction as a sampling task in which structures are generated independently, we formulated it as an ordered dependency between successive structural-state representations. We extended the folding-direction chain from sequence → profile → structure to sequence → profile → structure₁ → structure₂ (**Fig. 3b**). During fine-tuning, the model was trained to sequentially predict two experimentally observed conformations, thereby conditioning generation of the second structure on the first. This recomposition changes the dependency structure of generation while retaining the pretrained architecture, modalities and attention mechanisms, without introducing additional task-specific architectural components.

We fine-tuned ProteinReasoner on a filtered CoDNaS dataset^31^ containing 5,714 pairs of experimentally resolved conformations (see **Methods**), and evaluated it on two independent benchmarks: ligand-induced apo/holo transitions^32^ and fold-switching proteins^33^. Performance was assessed using two complementary metrics: TM-ens, which measures how well the predicted ensemble covers experimentally observed conformations, and ResFlex^27^, which quantifies agreement between predicted and observed residue-level flexibility patterns (**Extended Data Fig. 4a**). These complementary metrics capture global structural accuracy and local conformational variability.

Across both benchmarks, fine-tuned ProteinReasoner captured residue-level flexibility effectively, including the highest ResFlex on the fold-switching benchmark, while achieving TM-ens second only to AlphaFoldFlow among the compared methods (**Fig. 3c,d**). This performance, achieved after fine-tuning on substantially fewer conformations than the compared methods (**Fig. 3e**), shows that the modality chain can be reorganized for a distinct protein modelling task without introducing task-specific architectural components.

To isolate the contribution of cross-conformation conditioning, we compared consecutive generation with a three-modality baseline (sequence → profile → structure) in which conformations were sampled independently. Relative to the pretrained models, consecutive generation generally improved ResFlex while maintaining or improving TM-ens, whereas sampling produced little ResFlex gain on the apo/holo benchmark and, on the fold-switching benchmark, increased conformational variability at the expense of structural fidelity (**Fig. 3f**). Consecutive generation achieved higher TM-ens than sampling for both model scales on apo/holo (650M, 0.853, 95% CI 0.831–0.874 versus 0.836, 0.810–0.859; 3B, 0.861, 0.838–0.882 versus 0.847, 0.823–0.869) and fold-switching (650M, 0.682, 0.638–0.726 versus 0.663, 0.617–0.709; 3B, 0.706, 0.662–0.746 versus 0.675, 0.629– 0.717), with all paired comparisons significant (two-sided Wilcoxon tests, all P < 10⁻⁷). On apo/holo, consecutive generation also increased mean ResFlex for both scales, reaching significance for 650M (P = 0.021). These results indicate that cross-conformation conditioning directs fine-tuning towards better recovery of experimentally observed alternative states than independent sampling.

We next examined whether the added structural state influenced generation. Averaged across layers and heads, structure₂ directed 39.6% of its cross-modality attention to structure₁, more than to sequence (25.0%) or profile (9.4%) (**Extended Data Fig. 4b**), showing that the first structure was a major source of context during generation. We then compared two generation modes: conditioning the generation of structure₂ on a provided structure₁, and consecutively generating both structures without a provided conformation. Test proteins were stratified by the pretrained model’s intrinsic preference for either reference conformation, allowing the effect of conditioning to be separated from pre-existing generative bias. Across both benchmarks and all preference groups, providing structure₁ increased the proportion of structure₂ predictions closer to structure₁ relative to consecutive generation (**Extended Data Fig. 4c**). This effect persisted regardless of the pretrained model’s preferred conformation, supporting effective conditioning of second-conformation generation on the first structure.

As a representative case, we examined saccharopine reductase from *Magnaporthe grisea*, which adopts distinct apo and holo conformations associated with ligand binding^34^ (PDB IDs 1E5L and 1E5Q; **Fig. 3g**). Given the sequence and MSA, fine-tuned ProteinReasoner-3B generated 20 structures spanning conformations between the two experimental states, with progressive translation and rotation of domain I relative to domains II and III. Analysis of 180 additional predictions showed a continuous distribution across the RMSD range between the apo and holo structures, without discrete clusters or large gaps (**Fig. 3h**).

### 2.4 Experimental feedback as a modality for in-context protein optimization

Protein optimization seeks to improve biochemical properties such as stability, activity, specificity and binding affinity through sequence modification^6^. A common computational approach is to fine-tune a property predictor that ranks mutant sequences for experimental testing (**Fig. 4a**). Before introducing feedback-conditioned optimization, we established this conventional baseline by fine-tuning ProteinReasoner on the Megascale thermostability dataset^35^ using the same data splits as ThermoMPNN^36^ and ThermoMPNN-D^37^. Fine-tuned ProteinReasoner was competitive with ThermoMPNN on all-mutant and single-mutant evaluations, while both model scales achieved higher mean Spearman correlation and AUROC than ThermoMPNN-D on double mutants, although these differences were not significant (**Fig. 4b**). This establishes fine-tuned ProteinReasoner as a competitive baseline for evaluating the additional benefit of target-specific experimental feedback.

**Fig. 4.**
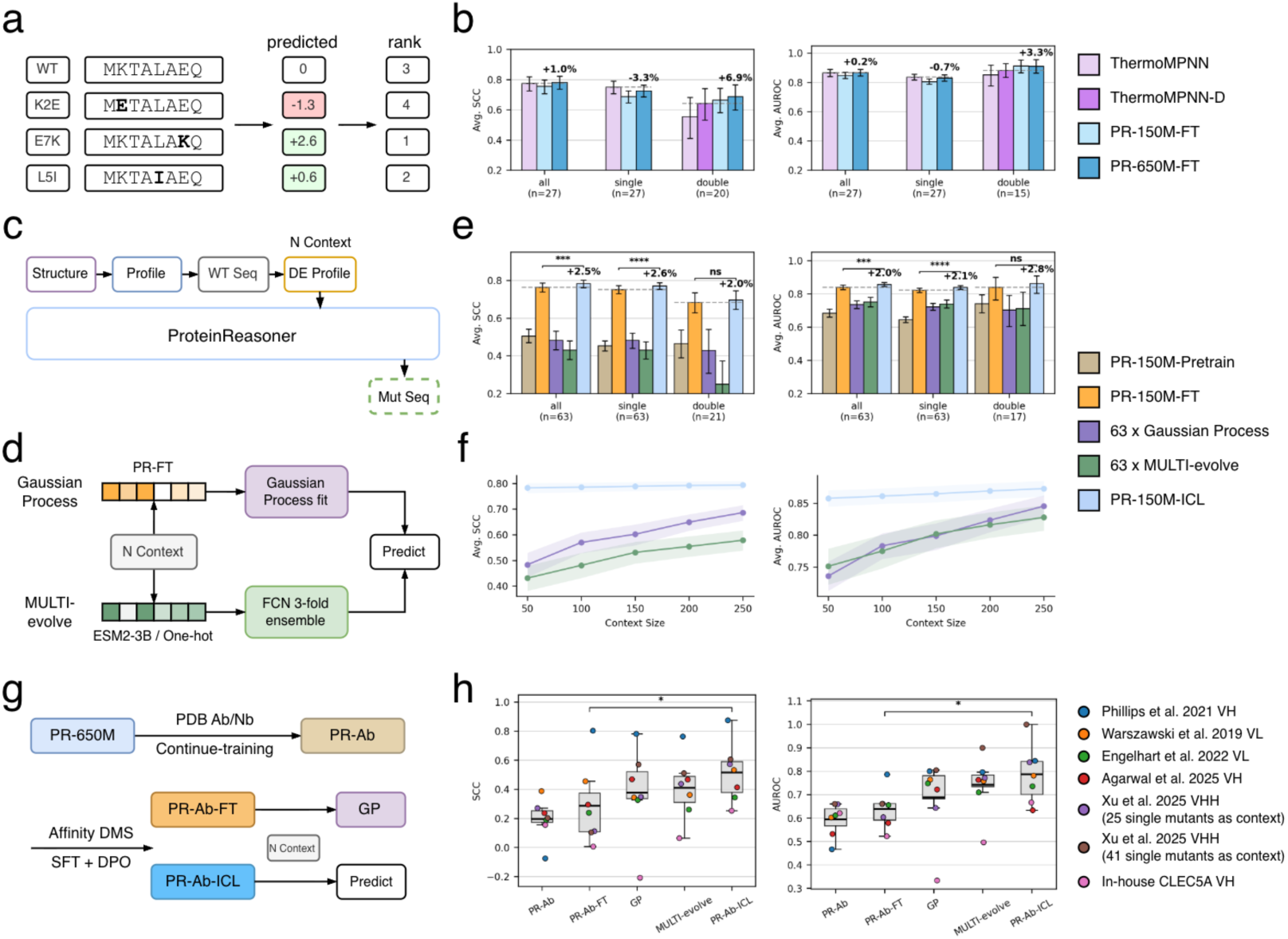
Modality-chain reasoning enables protein optimization from experimental feedback. **a**, Mutation-effect prediction as a ranking task for mutagenesis recommendation. **b**, Thermostability prediction on the Megascale dataset, evaluated by average per-protein Spearman correlation coefficient (SCC; left) and area under the receiver operating characteristic curve (AUROC; right) for all mutant sequences, single mutants and double mutants. Percentages indicate the relative difference between PR-650M-FT and the best-performing baseline. **c**, ProteinReasoner in-context learning (ICL) framework. A directed evolution (DE) profile constructed from N experimentally scored context mutant sequences extends the input modality chain and conditions the prediction of new mutant sequences without target-specific model updates. **d**, Target-specific Gaussian process (GP) and MULTI-evolve baselines trained using the same context mutant sequences. **e**, Performance on 63 held-out Megascale proteins using 50 context mutant sequences. Percentages indicate the relative improvement of PR-150M-ICL over PR-150M-FT. **f**, Megascale performance as a function of context size. **g**, Adaptation of ProteinReasoner to antibody and nanobody affinity prediction through continued pretraining and task-specific fine-tuning. **h**, SCC (left) and AUROC (right) across seven held-out antibody and nanobody evaluation cases. Coloured points denote individual datasets. In **b,e**, error bars indicate 95% bootstrap confidence intervals across proteins; in **f**, shaded regions indicate 95% bootstrap confidence intervals. Statistical significance in **e,h** was assessed using two-sided paired Wilcoxon signed-rank tests with unadjusted *P* values. ns, *P* ≥ 0.05; *, *P* < 0.05; **, *P* < 0.01; ***, *P* < 0.001; ****, *P* < 0.0001.

However, a fixed property predictor may not generalize across diverse proteins, particularly for target-specific objectives such as binding affinity. Protein optimization therefore often proceeds through iterative rounds of mutagenesis and experimental testing, with newly acquired measurements guiding subsequent designs. This iterative workflow embodies the lab-in-the-loop concept^38^ (**Extended Data Fig. 5a**), with active learning^39^ serving as a central strategy for selecting new mutant sequences and repeatedly updating a target-specific predictor, as exemplified by MULTI-evolve^40^ and EVOLVEpro^41^. Because each experimental round typically yields only a small number of labelled mutant sequences, these updates can be data-inefficient and sensitive to modelling choices.

We instead represent experimental feedback as an additional modality within an in-context learning (ICL)^42^ framework (**Fig. 4c**). Specifically, we extend the inverse-folding chain from structure → evolutionary profile → wild-type sequence to structure → evolutionary profile → wild-type sequence → directed evolution (DE) profile → mutant sequence. The DE profile summarizes experimentally characterized mutants as a position-wise representation of target-specific sequence–property relationships in the same matrix format as the evolutionary profile (see **Methods**). Following property-level adaptation on this extended chain, ProteinReasoner can condition on the encoded experimental feedback for mutant-sequence scoring or generation without fitting or updating parameters for each new target.

We evaluated this ICL formulation as a proof of concept on the Megascale dataset using a sequence-clustered split with 63 held-out test proteins (see **Methods**). Starting from ProteinReasoner-150M, we trained a single ICL model (PR-150M-ICL) using supervised fine-tuning (SFT) followed by direct preference optimization (DPO)^43^ on the extended chain containing the DE profile (**Extended Data Fig. 5b**). For each test protein, the DE profile was constructed from 50 experimentally scored mutant sequences, and performance was evaluated on the remaining mutant sequences. We compared PR-150M-ICL with pretrained ProteinReasoner and a matched no-context model (PR-150M-FT), trained using the same procedures but with the original structure → evolutionary profile → mutant sequence chain (**Extended Data Fig. 5c**). We also included two target-specific active-learning baselines: Gaussian process regression^44^ using PR-150M-FT embeddings and MULTI-evolve^40^. For each baseline, a separate model was fitted to each test protein using the same 50 mutant sequences supplied as context to PR-150M-ICL (**Fig. 4d**).

Across the 63 held-out proteins, PR-150M-ICL achieved higher mean Spearman correlation and AUROC than the matched PR-150M-FT baseline across all mutant sequences, single mutants and double mutants (**Fig. 4e**). The improvements were significant for the all-mutant and single-mutant evaluations in both metrics (two-sided paired Wilcoxon tests, all *P* < 0.001), while the smaller double-mutant subset showed the same directional advantage. Because the models shared the same pretrained foundation and SFT–DPO procedure, this comparison supports the contribution of target-specific feedback introduced through the DE-profile modality beyond property knowledge transferred across proteins.

With 50 context mutant sequences, PR-150M-FT also outperformed both active-learning baselines, despite receiving no target-specific feedback. This result highlights the strength of directly applying the transferred predictor when only limited measurements are available for a new target. As the context size increased from 50 to 250, both active-learning methods improved, but PR-150M-ICL maintained higher Spearman correlation and AUROC across all settings (**Fig. 4f**; **Extended Data Fig. 5d,e**). The context-scaling results therefore indicate that PR-150M-ICL combines transferable thermostability knowledge with increasing amounts of target-specific experimental information more effectively than fitting a separate model to the same measurements.

Megascale provides a useful proof of concept, but thermostability is relatively transferable across proteins. We therefore next evaluated whether the ICL framework extends to affinity maturation, a more target-specific setting in which beneficial mutations depend strongly on the antigen, scaffold and local sequence context.

For this evaluation, we adapted ProteinReasoner-650M to antibody and nanobody sequences through continued pretraining on 24,582 single-chain PDB entries, yielding PR-Ab. We then fine-tuned PR-Ab on 22 public affinity DMS datasets^45–48^ using the same SFT and DPO procedures as in the Megascale experiments, producing PR-Ab-FT and PR-Ab-ICL (**Fig. 4g**; **Extended Data Fig. 6a**). We evaluated seven additional test cases^45–49^ spanning different levels of difficulty, defined by binding-target exposure during training and CDR sequence identity to the training data (**Extended Data Fig. 6b and 7**). For each case, 25–56 experimentally scored context mutant sequences were used to construct the DE profile or fit the target-specific baselines, and performance was evaluated on up to 512 additional mutant sequences whose mutated positions were represented in the context set. Comparisons included PR-Ab, PR-Ab-FT, Gaussian process regression using PR-Ab-FT embeddings, and MULTI-evolve.

Across the seven affinity-maturation cases, PR-Ab-ICL achieved higher Spearman correlation and AUROC than PR-Ab-FT, with mean improvements of 0.227 and 0.147, respectively (**Fig. 4h**; **Extended Data Fig. 6b**). Both differences were significant across paired cases (two-sided paired Wilcoxon tests, both *P* = 0.016), including gains on binding targets not represented in the training datasets. These results show that the extended modality chain enables target-specific adaptation from experimental context beyond the affinity knowledge transferred during fine-tuning.

Relative to the target-specific active-learning baselines, PR-Ab-ICL performed best in five cases, comparably to Gaussian process regression in one and worse in one, and achieved the highest mean Spearman correlation and AUROC across the seven cases (**Fig. 4h; Extended Data Fig. 6b**). Given the limited availability of suitable affinity-maturation datasets, these cases provide a focused rather than comprehensive evaluation. Across thermostability and affinity maturation, these results establish an in-context learning paradigm in which a shared foundation model combines transferable property knowledge with target-specific experimental feedback without further parameter updates.

## 3. Discussion

Here we formulate reasoning in scientific foundation models as sequential modelling across explicit, interdependent representations, in which each representation constrains the prediction or generation of the next. ProteinReasoner provides an operational realization of this idea: controlled comparisons support a functional role for informative intermediate representations, while reorganizing or augmenting the chain allows the same pretrained backbone to accommodate new dependencies. These results provide an empirically supported formulation of reasoning across interdependent scientific representations.

Unlike textual reasoning in large language models, modality-chain reasoning operates over domain-specific scientific representations rather than verbal rationales. In ProteinReasoner, these include quantitative evolutionary profiles, discrete structural states and experimentally derived representations of sequence–property relationships. Keeping these representations explicit within a shared autoregressive context allows their dependencies to be specified and examined: intermediate profiles can be supplied, inferred or replaced, and later outputs can be tested for dependence on preceding representations. This organization also allows modality chains to be reversed, reorganized or augmented with task-relevant representations while retaining the same pretrained backbone. Modality-chain reasoning therefore connects a concrete formulation of scientific reasoning with a unified strategy for adapting foundation models across protein tasks.

A particularly consequential extension is the incorporation of experimental feedback through in-context learning. Conventional active-learning approaches use newly acquired measurements to fit or update a target-specific predictor. ProteinReasoner instead summarizes experimentally characterized mutants as a DE profile, making target-specific sequence–property relationships an explicit representation within the modality chain. This allows the shared model to combine property knowledge transferred across proteins with information specific to the current optimization case, without further parameter updates. In both thermostability and affinity maturation, this formulation improved over matched fine-tuned models and achieved stronger overall performance than target-specific active-learning baselines. These results show that experimental feedback can serve directly as context for prediction and generation within a shared foundation model, rather than solely as supervision for retraining.

Several limitations remain. First, multimodal pretraining was constrained by the computational cost of MSA search and profile construction, leaving the scaling behaviour of modality-chain reasoning insufficiently characterized. Moreover, multimodal pretraining excluded proteins with fewer than ten aligned sequences, and performance when evolutionary information is sparse or unavailable remains to be evaluated. Second, the current framework is limited in the scope of structural systems represented within a modality chain. Multiple-conformation prediction currently models only two structural states, and ProteinReasoner operates on individual protein chains. In the affinity-maturation experiments, for example, the antigen is not explicitly represented. Extending modality chains with molecular dynamics trajectories or larger conformational ensembles could enable reasoning over more continuous structural transitions. Incorporating interacting chains or complex-level representations could further support direct modelling of molecular interactions. Third, the in-context learning framework currently requires property-level adaptation before it can interpret experimental feedback in context, and the protein-optimization experiments in this study were evaluated retrospectively using existing experimental measurements. Future work should extend this framework to iterative and multi-objective optimization and evaluate its ability to guide successive design–test cycles prospectively in laboratory workflows. More broadly, extending modality-chain reasoning to additional scientific representations and domains will allow the scientific community to test, refine and expand this strategy across diverse problems.

## Methods

### ProteinReasoner architecture and modality representations

ProteinReasoner is a decoder-only Transformer that represents heterogeneous protein information as ordered modality segments within a shared autoregressive context. During pretraining, the model processes two modality chains: sequence → evolutionary profile → structure and structure → evolutionary profile → sequence. Each modality is delimited by modality-specific beginning- and end-of-modality tokens, allowing the model to preserve modality identity while learning dependencies across successive segments.

Amino acid sequences are tokenized at residue resolution using a 26-token amino acid vocabulary plus 11 special tokens. Protein structures are represented by residue-level atomic coordinates and discretized using the DPLM-2 structure tokenizer, with a vocabulary of 8,192 structure tokens. We used the *airkingbd/struct_tokenizer* checkpoint from Hugging Face. Evolutionary information is represented as a position-wise profile derived from a multiple sequence alignment (MSA). Given n aligned sequences of length L, the evolutionary profile *P* ∈ ℝ*^L^*^×21^ is defined as

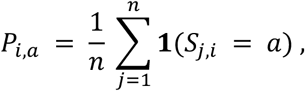

where *S*_j,*i*_ denotes the residue at position *i* in aligned sequence *j*, *a* ranges over the 20 standard amino acids and the gap character, and **1**(⋅) is the indicator function. Each row therefore forms a categorical distribution describing residue preferences at one sequence position.

Sequence and structure tokens are mapped through modality-specific embedding layers, whereas evolutionary profiles are projected through a linear layer. Each residue-aligned modality initially yields *L* embeddings in ℝ*^L^*^×*D*^. After prepending and appending modality-specific boundary-token embeddings, each modality segment has length *L* + 2 and is represented in ℝ^(*L*+2)×*D*^, where *D* is the Transformer hidden dimension. The sequence embedding and output layers are initialized from the sequence-only model obtained during the first pretraining stage; the profile- and structure-specific input and output modules are initialized from scratch.

The modality embeddings are concatenated and processed by *N* causal Transformer decoder layers. Rotary positional encodings are applied using residue indices that restart for each modality, aligning corresponding residue positions across modality segments. Outputs from the final Transformer layer are separated by modality and passed through modality-specific MLP prediction heads for amino acid tokens, evolutionary-profile distributions and structure tokens. Model dimensions and architecture specifications are provided in **Supplementary Table 2**.

### Pretraining data and optimization

ProteinReasoner was trained in two stages. The first stage used the sequence-only protein language model described in the previous study^14^, trained on approximately one trillion amino-acid tokens, to initialize the sequence embedding, Transformer backbone and sequence output head.

For multimodal pretraining, we constructed a dataset comprising 9,450,396 proteins from the AlphaFold Protein Structure Database and 279,921 experimentally resolved chains from PDB-REDO. AlphaFoldDB proteins were sampled from 2.3 million non-singleton Foldseek^50^ clusters, and entries with mean pLDDT below 70 were excluded. Proteins appearing in any downstream test set were removed from the pretraining data. For each retained protein, homologous sequences were searched against UniRef30 using HHblits^51^ v3.3.0 with -n 2 -cov 70 -id 90 -diff 2000; proteins with fewer than ten aligned sequences were excluded.

ProteinReasoner was trained autoregressively on the sequence → evolutionary profile → structure and structure → evolutionary profile → sequence chains. For sequence and structure modalities, the model predicts the next discrete token using cross-entropy loss. For the evolutionary-profile modality, it predicts the position-wise amino-acid distribution using Kullback–Leibler divergence. For an input chain *X*, the total loss is

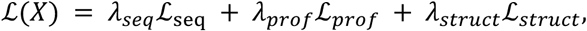

where the modality-specific weights were set separately for each model scale and are reported in

### Supplementary Table 3

Models were trained for 60,000 optimization steps with a global batch size of 1,024 and a maximum sequence length of 3,072, corresponding to up to 1.89 × 10^11^ multimodal training tokens. Training used the Adam optimizer with *β*_1_ = 0.9, *β*_2_ = 0.95, with weight decay 0.1, dropout 0.1 and bfloat16 mixed precision. The learning rate was increased linearly during warm-up and then decayed using a cosine schedule. Model-specific learning rates, warm-up proportions and loss weights are provided in **Supplementary Table 3**.

### Zero-shot benchmark datasets and evaluation

#### Structure prediction and inverse folding datasets

We evaluated ProteinReasoner on 110 single-chain proteins from CAMEO released between 1 July 2021 and 1 June 2022, 43 CASP14 targets and 51 CASP15 targets. Experimentally resolved structures were obtained from the Protein Data Bank, and MSAs were generated against UniRef30 using MMSeqs2^52^ following the ColabFold^53^ search procedure. We additionally evaluated 449 single-chain proteins from the MultiFlow^54^ PDB holdout set. For inverse folding, 393 proteins were retained after excluding cases in which the reference sequence could not be matched to the sequence extracted from the corresponding PDB structure. All benchmark proteins were removed from the ProteinReasoner pretraining data. To assess generalization beyond close homologues, the CASP, CAMEO and PDB holdout sets were pooled and filtered to retain proteins sharing less than 50% or 30% sequence identity with any ProteinReasoner pretraining sequence.

#### Baseline models

We evaluated ESM3-Open-1.4B using the *esm3-sm-open-v1* checkpoint and the official ESM inference procedures. For structure prediction, the amino acid sequence track was provided and the structure track was generated; for inverse folding, atomic coordinates were tokenized and used to generate the sequence track. We evaluated generation with *num_steps = 1* and *num_steps = L*, where *L* is the protein length, and reported the better-performing setting for each task. DPLM-2-650M and DPLM-2-3B were evaluated using the *airkingbd/dplm2_650m* and *airkingbd/dplm2_3b* checkpoints and the official *generate_dplm2.py* script with default settings, using argmax decoding for 100 iterations.

#### Structure prediction

Predicted structure tokens were decoded into atomic coordinates using the corresponding structure decoder. Predicted and experimental structures were aligned using US-align v20220924 with -outfmt 2, from which root-mean-square deviation and TM-score were calculated.

#### Inverse folding

Sequence recovery was measured by amino acid recovery,

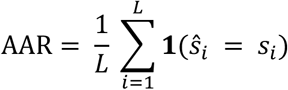

where *s_i_* and *ŝ_i_* denote the native and generated amino acids at position *i*, respectively. Structural consistency was evaluated using self-consistency TM-score. For each evaluated protein, the generated sequence was folded using AlphaFold2 and compared with the input structure using US-align.

#### ProteinGym fitness prediction

We evaluated 217 assays from the ProteinGym DMS substitution benchmark. Wild-type sequences, structures, MSAs and experimental measurements were obtained from the official ProteinGym release. Evolutionary profiles were computed using RSALOR^55^ without incorporating relative solvent accessibility. Given profile-head logits 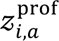 and sequence-head logits 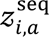 for amino acid *a* at position *i*, we defined the combined log-probability as

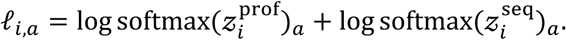

For a mutation from wild-type residue 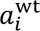 to mutant residue 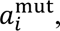, the predicted effect was

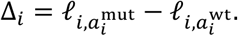

For sequences containing multiple substitutions, the mutation scores were summed across mutated positions. Prediction files were evaluated using the official ProteinGym pipeline, and performance was reported as Spearman correlation with the experimental measurements.

#### Statistical analysis

For structure-prediction and inverse-folding benchmarks, mean performance and 95% confidence intervals were estimated across proteins by non-parametric bootstrap resampling with replacement (10,000 resamples). Statistical comparisons between models used two-sided paired Wilcoxon signed-rank tests, with proteins as the paired evaluation units. Only proteins with finite measurements for both models were included in each comparison, and missing measurements were not imputed. Statistical tests were performed separately for each benchmark and metric, and reported *P* values were unadjusted. Associations between profile KLD and downstream performance were assessed using two-sided Spearman rank correlation. Linear fits were restricted to proteins with KLD ≤ 2.

### Multiple-conformation prediction

ProteinReasoner-650M and ProteinReasoner-3B were adapted for multiple-conformation prediction by extending the folding chain to sequence → evolutionary profile → structure₁ → structure₂. The additional structure segment used the same tokenizer, input embedding and output head as the original structure modality; all other architectural components remained unchanged. Models were fine-tuned on paired experimentally resolved conformations using cross-entropy supervision over the two structure-token segments. If *I_s_*_1_ and *I_s_*_2_ denote the residue-token positions of structure₁ and structure₂, respectively, the fine-tuning objective was

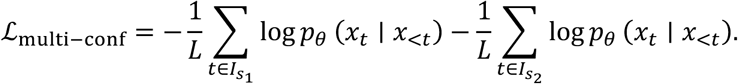

Training hyperparameters are provided in **Supplementary Table 4**.

Paired conformations were obtained from CoDNaS, yielding an initial set of 45,363 pairs. We retained pairs with RMSD between 0.1 and 10.0 Å, sequence identity greater than 90%, and protein lengths between 50 and 1,024 residues. To avoid leakage into the evaluation benchmarks, retained sequences were additionally required to share less than 30% sequence identity with all test proteins. The resulting dataset comprised 5,714 conformation pairs, divided into 5,241 training and 473 validation pairs. MSAs were generated against UniRef30 using MMseqs2.

Evaluation was performed on the apo/holo and fold-switching benchmark sets distributed through the EigenFold repository. For each protein, the sequence provided in the benchmark metadata was used as input, and an MSA was generated against UniRef30 using MMseqs2. Each model generated an ensemble of conformations, which was evaluated using TM-ens and ResFlex.

TM-ens measures coverage of the experimentally observed conformations. Given a predicted ensemble *P* and a reference set ℛ, it is defined as

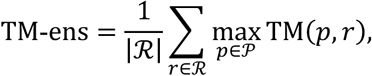

where TM(*p*, *r*) is the TM-score between predicted structure *p* and reference conformation *r*.

ResFlex measures agreement between predicted and experimentally observed residue-level conformational variability. Structures within each ensemble were aligned, residue-wise flexibility was calculated from the displacement of centred Cα coordinates, and the Spearman correlation between the predicted and reference flexibility profiles was reported.

Multiple-conformation prediction was evaluated following the ESMDiff protocol (https://github.com/lujiarui/esmdiff/blob/main/analysis/apo_analysis.py). For the published baselines, five structures were sampled per target; ResFlex was computed from the 10 pairwise comparisons among these structures, and TM-ens was calculated by independently selecting the best-matching structure for each of the two reference conformations. Because ProteinReasoner generates conformations in pairs, ResFlex was computed from 10 within-pair comparisons across 10 generated pairs. For TM-ens, the first five generated pairs were used, with the two structures in each pair assigned exclusively to the two reference conformations to maximize their summed TM-score, yielding five candidate scores per reference conformation. Results for AlphaFlow, ESMFlow and ESMDiff were obtained from the ESMDiff study, which used the same benchmark sets and evaluation procedures. For the ESMDiff model family, the best reported variant was selected separately for each metric. Model-training set sizes were taken from the corresponding publications. Target-level values were not available for uncertainty estimation.

For the ablation analysis, mean ResFlex and TM-ens values and 95% confidence intervals were estimated across proteins by non-parametric bootstrap resampling with replacement (10,000 resamples). Consecutive generation and independent sampling were compared using two-sided paired Wilcoxon signed-rank tests, with proteins as the paired evaluation units. Only proteins with finite measurements for both models were included.

### Protein optimization models and training

ProteinReasoner was adapted for protein optimization under conventional fine-tuning (FT) and in-context learning (ICL) settings. The FT model used the three-modality chain structure → evolutionary profile → mutant sequence. The ICL model extended this chain to structure → evolutionary profile → wild-type sequence → directed evolution (DE) profile → mutant sequence. The wild-type structure and evolutionary profile were used in both settings.

The DE profile represents experimental feedback from *K* characterized mutant sequences as a position-wise distribution *D* ∈ ℝ*^L^*^×20^ over the standard amino acids. Given context sequences 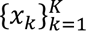 and their wild-type-centred, *z*-score-normalized property values 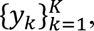, sequence weights were estimated offline using ordinary kriging with a Hamming-similarity kernel and projected back to residue space. A position-wise softmax was then applied so that *D_i_*_,*a*_ ≥ 0 and ∑*_a_ D_i_*_,*a*_ = 1. The resulting tensor was supplied to ProteinReasoner during both training and inference; it was not generated from the context measurements by the model. The complete construction and its equivalence to a Gaussian-process posterior projected into residue space are provided in the **Supplementary Information**.

Before fine-tuning, the DE-profile input projection was initialized from the pretrained evolutionary-profile projection by removing the weights associated with the gap input dimension, and the mutant-sequence output head from the pretrained sequence head. For affinity maturation, we further introduced a lightweight, profile-conditioned causal cross-attention module between the Transformer backbone and mutant-sequence decoder. During DPO of the affinity-maturation ICL model, the mutant-sequence distribution was further fused with the input DE profile through a case-level gated residual in amino-acid log-probability space; the gate was predicted from the DE-profile hidden states and shared by both sequences in each preference pair. Full architectural details are provided in the **Supplementary Information**.

For affinity maturation, ProteinReasoner-650M was first continued for 300 optimization steps on 24,582 single-chain antibody and nanobody entries from the Protein Data Bank released by 29 May 2024, using the original pretraining chains and objective, yielding the PR-Ab checkpoint. VH, VL and VHH sequences were paired with their experimental structures, and evolutionary profiles were generated by searching UniRef30, filtering homologues at a minimum sequence identity of 0.2 and an E-value threshold of 1*e*^-3^.

During supervised fine-tuning (SFT), mutant sequences with experimentally measured scores greater than that of the corresponding wild type were used as positive examples. Supervision was applied only to residue tokens in the mutant-sequence segment. For a mutant sequence *x*^mut^ of length *L*, the sequence loss was

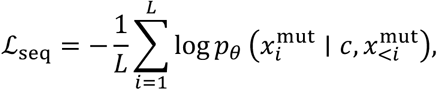

where *c* denotes the preceding modality context. For ICL models, we additionally supervised reconstruction of the supplied DE profile using

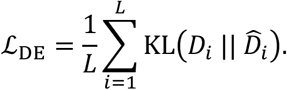

The total ICL supervised objective was

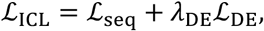

with *λ*_DE_ = 0.1.

The supervised models were subsequently aligned using direct preference optimization. Preference pairs comprised two mutant sequences from the same wild-type protein, with the higher-scoring sequence designated as the preferred sequence. Given preferred and less-preferred sequences *x*^+^ and *x*^-^, the DPO loss was

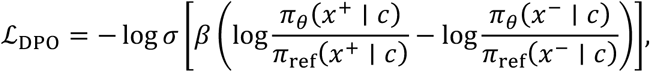

where *π*_ref_ is the corresponding supervised fine-tuned model and *β* controls the strength of preference optimization. Within each experiment, FT and ICL models used matched training settings wherever possible; the Megascale FT model was selected by early stopping because this improved validation performance. Full SFT and DPO hyperparameters and training-example construction procedures are provided in **Supplementary Tables 5–7** and **Algorithms 1–2**.

### Protein optimization datasets and evaluation

#### Thermostability fine-tuning benchmark

To compare ProteinReasoner with ThermoMPNN and ThermoMPNN-D, we fine-tuned ProteinReasoner-150M and ProteinReasoner-650M on the Megascale thermostability dataset using the same protein-level split adopted by both baselines, comprising 238 training, 31 validation and 27 test proteins. ThermoMPNN and ThermoMPNN-D were evaluated using their official implementations and checkpoints on the same test set. Performance was assessed separately across all mutant sequences, single mutants and double mutants.

#### Megascale ICL evaluation

For the in-context learning experiments, Megascale proteins were clustered by wild-type sequence similarity and divided into 411 training, 4 validation and 63 held-out test proteins. Protein structures were predicted using AlphaFold2, and MSAs were generated against UniRef30 using MMseqs2. For 159 of the 478 proteins without sufficient MSA coverage, evolutionary profiles were generated using ProteinReasoner-650M. Experimental measurements were centred on the corresponding wild-type score, such that positive values indicated stabilizing mutations.

For each test protein, mutant sequences were ranked by sequence log-likelihood under pretrained ProteinReasoner. The top *K* sequences were used to construct the DE profile for PR-150M-ICL and as the target-specific training set for the active-learning baselines; all remaining mutant sequences were reserved for evaluation. The primary comparison used *K* = 50, and the context-scaling analysis varied *K* ∈ {50, 100, 150, 200, 250}. Thus, all context-dependent methods received the same experimentally characterized mutant sequences.

#### Affinity-maturation datasets

We assembled 28 antibody and nanobody affinity datasets from FLAb2, AbBiBench, BindingGYM, AlphaBind, a published LGR5-targeting VHH affinity-maturation study and an inhouse CLEC5A-targeting dataset. Twenty-two datasets were used for training, and seven held-out evaluation cases were constructed from the remaining datasets, including two different context–test partitions of the LGR5 dataset. Evaluation cases were separated from the training data by binding-target identity and CDR sequence similarity. CDR1, CDR2 and CDR3 were identified using ANARCI, concatenated and aligned to calculate pairwise identity between datasets.

Experimentally determined structures were used when available; otherwise, structures were predicted using AlphaFold2. Evolutionary profiles were generated by searching each VH, VL or VHH sequence against UniRef30, retaining homologues with at least 20% sequence identity and an E-value below 1*e*^-3^, and applying the standard ProteinReasoner profile-generation pipeline. Experimental measurements were directionally harmonized so that larger values indicated improved binding. Mutant sequences scoring above the corresponding wild type were treated as positives, and ties were excluded.

Four evaluation cases were derived from high-throughput mutational-scanning datasets. For each replicate, 50 single- and double-mutant sequences were sampled as context, with approximately 25% positives where available. Up to 512 remaining mutant sequences were evaluated, restricted to sequences whose mutated positions were represented in the context set. This procedure was repeated 16 times per case. For the first LGR5 case, 25 single mutants were randomly sampled as context, with the remaining 46 sequences used for evaluation. This procedure was repeated across 16 replicates. For the second LGR5 case, all 41 single mutants were used as context and 30 multi-mutant sequences as test data. For CLEC5A, 56 sequences containing one to three mutations were used as context and 27 sequences containing four to six mutations as test data. Complete dataset compositions and processed context–test splits are provided in the **Supplementary Data**.

#### Target-specific baselines

Gaussian-process regression and MULTI-evolve were fitted independently for each test protein using the same context mutant sequences supplied to the ICL model. For Gaussian-process regression, residue-level representations from the corresponding fine-tuned ProteinReasoner model were mean-pooled to obtain fixed-dimensional mutant-sequence embeddings. An exact Gaussian process with a constant mean, linear covariance kernel and Gaussian likelihood was optimized by maximizing the exact marginal log-likelihood, and candidates were ranked by posterior predictive mean.

MULTI-evolve was implemented using its official repository. We evaluated both flattened position-wise one-hot sequence representations and mean-pooled residue embeddings from ESM-2-3B. For each target protein and input representation, we evaluated 18 network configurations using three-fold cross-validation. The input representation and network configuration with the lowest mean validation mean-squared error were selected, and final predictions were obtained by averaging predictions from the corresponding three fold-specific models. Full baseline configurations are provided in the **Supplementary Information**.

#### Evaluation metrics

Performance was calculated separately within each wild-type protein or affinity-maturation case using Spearman correlation between predicted and experimental scores and the area under the receiver operating characteristic curve (AUROC). Megascale results were summarized by the mean across proteins. For affinity-maturation cases with repeated context sampling, metrics were first calculated for each replicate and then averaged within each case; these seven case-level values were used for across-case comparisons.

#### Statistical analysis

For the Megascale analyses, mean performance and 95% confidence intervals were estimated across proteins by non-parametric bootstrap resampling with replacement (10,000 resamples). Statistical comparisons used two-sided paired Wilcoxon signed-rank tests on proteins with finite measurements for both models. For affinity maturation, repeated context samplings were averaged within each case before paired comparisons across the seven evaluation cases. For Extended Data Fig. 6b, 95% confidence intervals for cases with 16 context-sampling repeats were estimated by bootstrap resampling across repeats. Reported *P* values were unadjusted.

### Additional analyses

#### Cross-modality attention analysis

To examine how ProteinReasoner used information from preceding modalities, we extracted self-attention weights from all Transformer layers and heads during inference. Tokens were grouped according to their modality segments. For each sample, attention from a source modality *S* to a preceding target modality *T* was calculated as

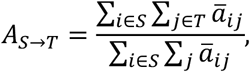

where *a*u*_i_*_j_ denotes the attention weight from query token *i* to key token *j*, averaged across layers and heads. Boundary tokens were excluded. Sample-level values were summarized across each benchmark dataset. For the profile-quality analysis, individual heads with high target-to-profile attention were further examined in relation to MSA depth or evolutionary-profile Kullback–Leibler divergence.

#### Conditional generation analysis

To assess whether structure₁ influenced generation of structure₂, we performed conditioned and consecutive generation on the apo/holo and fold-switching benchmarks. In the conditioned setting, one experimentally observed conformation was supplied as structure₁, and the fine-tuned model generated structure₂. In the consecutive setting, both structural states were generated without a provided conformation. Each prediction was assigned to the closer of the two reference conformations according to RMSD. To account for the pretrained model’s intrinsic preference for either reference state, proteins were stratified by a similarity vote obtained from five structures generated by pretrained ProteinReasoner-3B. The fine-tuned model then generated 20 structures per protein under each setting, and we calculated the proportion assigned to each reference conformation within every preference group.

#### Conformation-space analysis

We analysed saccharopine reductase from *Magnaporthe grisea* using the experimentally observed apo and holo structures (PDB IDs 1E5L and 1E5Q). Fine-tuned ProteinReasoner-3B generated 200 structures from the apo-state sequence and MSA. Each prediction was aligned to the reference structures using domain II and most of domain III, and Cα RMSD was calculated over domain I and the ligand-contacting loop in domain III (residues 2–121, 172–183 and 391–444). Predictions were positioned in a two-dimensional space according to their RMSD from the apo and holo references, and a density estimate was used to visualize their distribution between the two experimentally observed states.

## Data availability

Publicly available datasets used in this study were obtained from the AlphaFold Protein Structure Database, PDB-REDO, UniRef30, the Protein Data Bank, CAMEO, CASP14, CASP15, ProteinGym, Megascale, FLAb2, AbBiBench, BindingGYM and AlphaBind, as well as benchmark resources distributed with MultiFlow and EigenFold, as described in the Methods and cited in the corresponding publications. The CoDNaS conformation-pair dataset was provided directly by the CoDNaS authors.

## Acknowledgements

We thank Gustavo Parisi for assistance in accessing the CoDNaS dataset. We also thank Xing Rao, Sikai Zhang and Guoliang Li for support with large-scale MSA generation and for the internal AlphaFold2 prediction service used in this study.

## Author Contributions

C.L., L.C., and X.Z. conceived the research. M.Y., S.J., H.W., Z.G., C.L., and L.C. built the pretraining infrastructure. C.L. and L.C. performed model pretraining, fine-tuning and evaluation. C.L., L.C., and Y.G. performed dataset preprocessing and curation. J.Z. implemented the attention map analysis. D.H. conducted the multiple-conformation prediction case study. G.Z. fine-tuned the thermostability model used for comparison with ThermoMPNN. T.J. implemented the direct preference optimization algorithm. J.P. contributed to the design and evaluation of the affinity-maturation task. S.L. and Y.G. contributed to the conceptual development of the study and interpretation of the results. X.Z. and L.C. supervised the research. C.L. wrote the initial draft of the manuscript. All authors reviewed and edited the manuscript.

---

**Extended Data Fig. 1.**
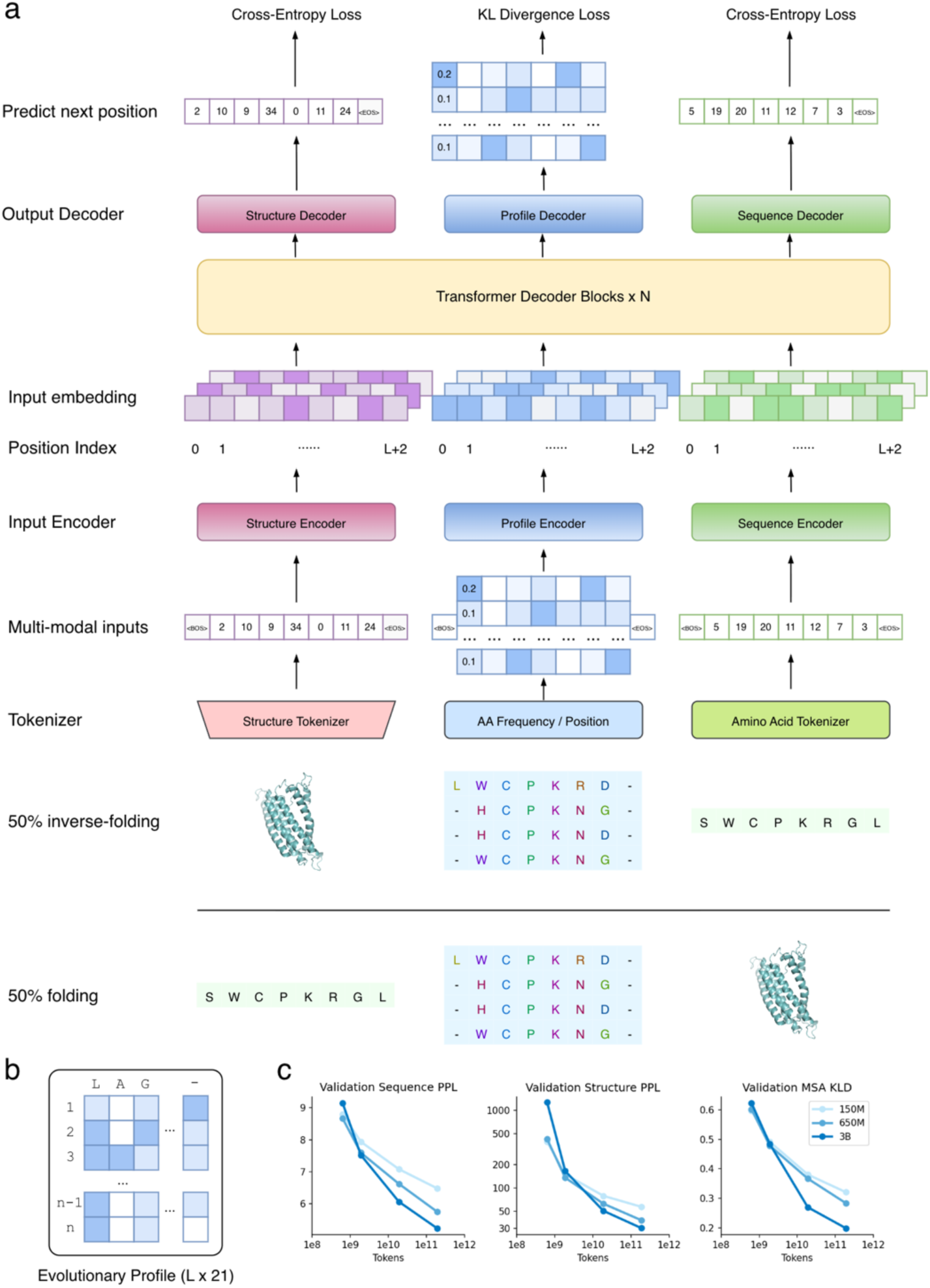
ProteinReasoner architecture and pretraining curves. **a**, Detailed architecture of ProteinReasoner. Multi-modal inputs are represented using modality-specific input modules: sequence and structure are tokenized, whereas the evolutionary profile is projected from its continuous position-wise representation. The resulting embeddings are concatenated and processed through shared Transformer blocks. Separate decoders predict sequence tokens, evolutionary profiles, and structure tokens, optimized using cross-entropy loss for sequence and structure and Kullback–Leibler divergence loss for the evolutionary profile. Pretraining uses equal proportions of folding and inverse-folding modality chains. **b**, Representation of the evolutionary profile as an *L* × 21 matrix, encoding position-specific amino acid frequencies including a gap character. **c**, Pretraining learning curves on held-out validation data. Shown are validation sequence perplexity, validation structure perplexity, and validation evolutionary profile KLD as a function of training tokens for the 150M, 650M, and 3B ProteinReasoner models.

**Extended Data Fig. 2.**
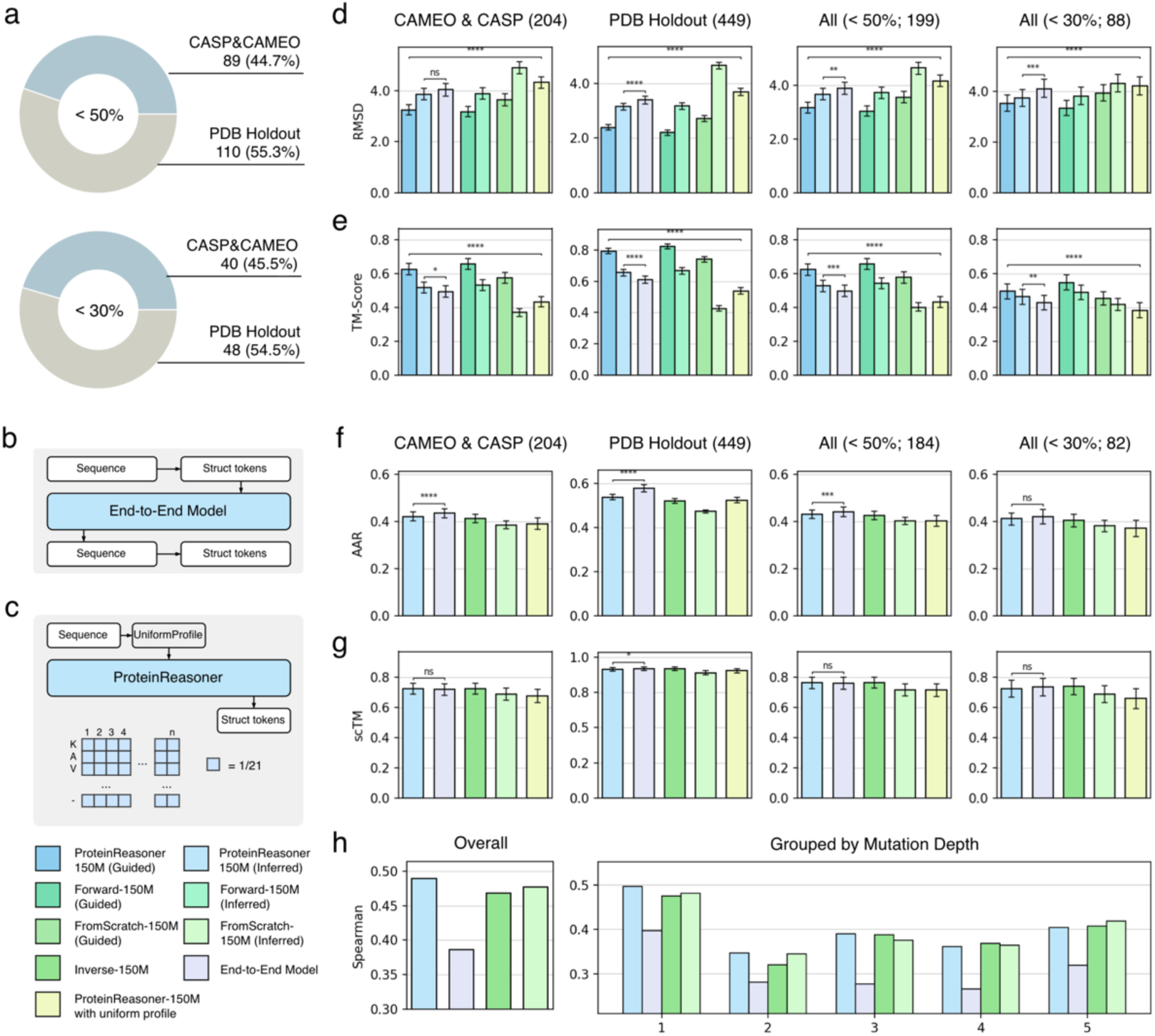
Ablation studies of ProteinReasoner design choices. **a**, Composition of the sequence similarity-controlled benchmark subsets, showing contributions from the combined CAMEO and CASP set and the PDB holdout set at sequence identity thresholds of <50% and <30% relative to the pretraining data. **b**, End-to-end pretraining baseline. Sequence and structure tokens are concatenated and modelled directly without an intermediate evolutionary profile. **c**, Uniform-profile ablation. ProteinReasoner is provided with a non-informative uniform evolutionary profile (1/21 for all amino acids and gap) instead of a profile derived from evolutionary signals. **d,e**, Structure prediction performance measured by RMSD (**d**) and TM-score (**e**) on the combined CAMEO and CASP set, the PDB holdout set, and the <50% and <30% sequence-identity subsets. **f,g**, Inverse folding performance measured by amino acid recovery (AAR; **f**) and self-consistency TM-score (scTM; **g**) on the corresponding benchmark sets. **h**, Zero-shot fitness prediction on ProteinGym, shown as the overall Spearman correlation (left) and stratified by mutation depth (right). In **d–g**, bars show mean performance and error bars indicate 95% bootstrap confidence intervals across proteins. Statistical brackets in **d,e** compare internally inferred ProteinReasoner-150M with the end-to-end ablation and guided ProteinReasoner-150M with the uniform-profile ablation; those in **f,g** compare internally inferred ProteinReasoner-150M with the end-to-end ablation. Statistical significance was assessed using two-sided paired Wilcoxon signed-rank tests. *P* values are unadjusted. ns, *P* ≥ 0.05; *, *P* < 0.05; **, *P* < 0.01; ***, *P* < 0.001; ****, *P* < 0.0001.

**Extended Data Fig. 3.**
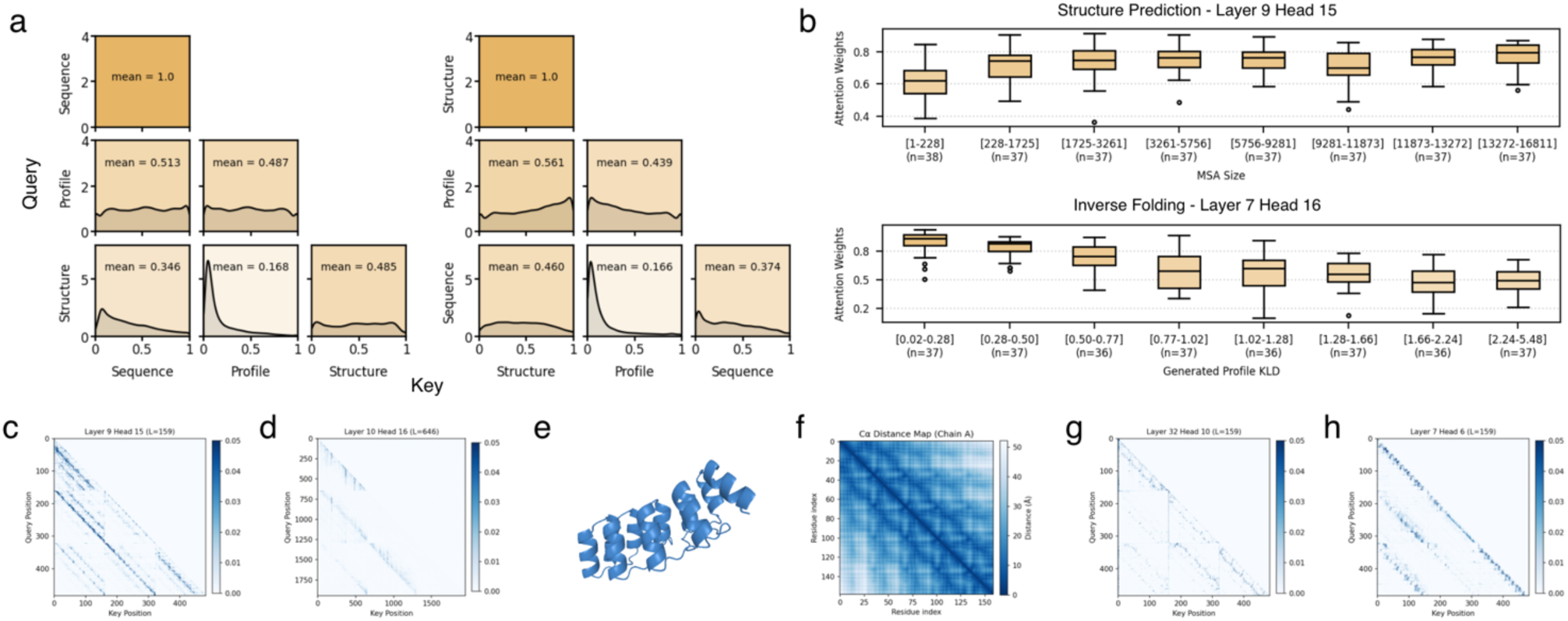
Cross-modality attention patterns in ProteinReasoner. **a**, Distributions of cross-modality attention weights in the folding (left) and inverse-folding (right) directions. Values indicate mean attention weights averaged across layers and heads. **b**, Attention weights of representative heads stratified by MSA depth for structure prediction (top) and inferred-profile Kullback–Leibler divergence (KLD) for inverse folding (bottom). **c,d**, Attention maps from representative profile-focused heads in the folding direction for protein 7P3I chain B (**c**) and in the inverse-folding direction for protein 7ESH chain A (**d**). **e,f**, Three-dimensional structure of 7P3I chain B (**e**) and its Cα distance map (**f**). **g,h**, Attention maps from additional representative heads in the folding (**g**) and inverse-folding (**h**) directions for 7P3I chain B, showing patterns that resemble residue–residue proximity and task-dependent variation.

**Extended Data Fig. 4.**
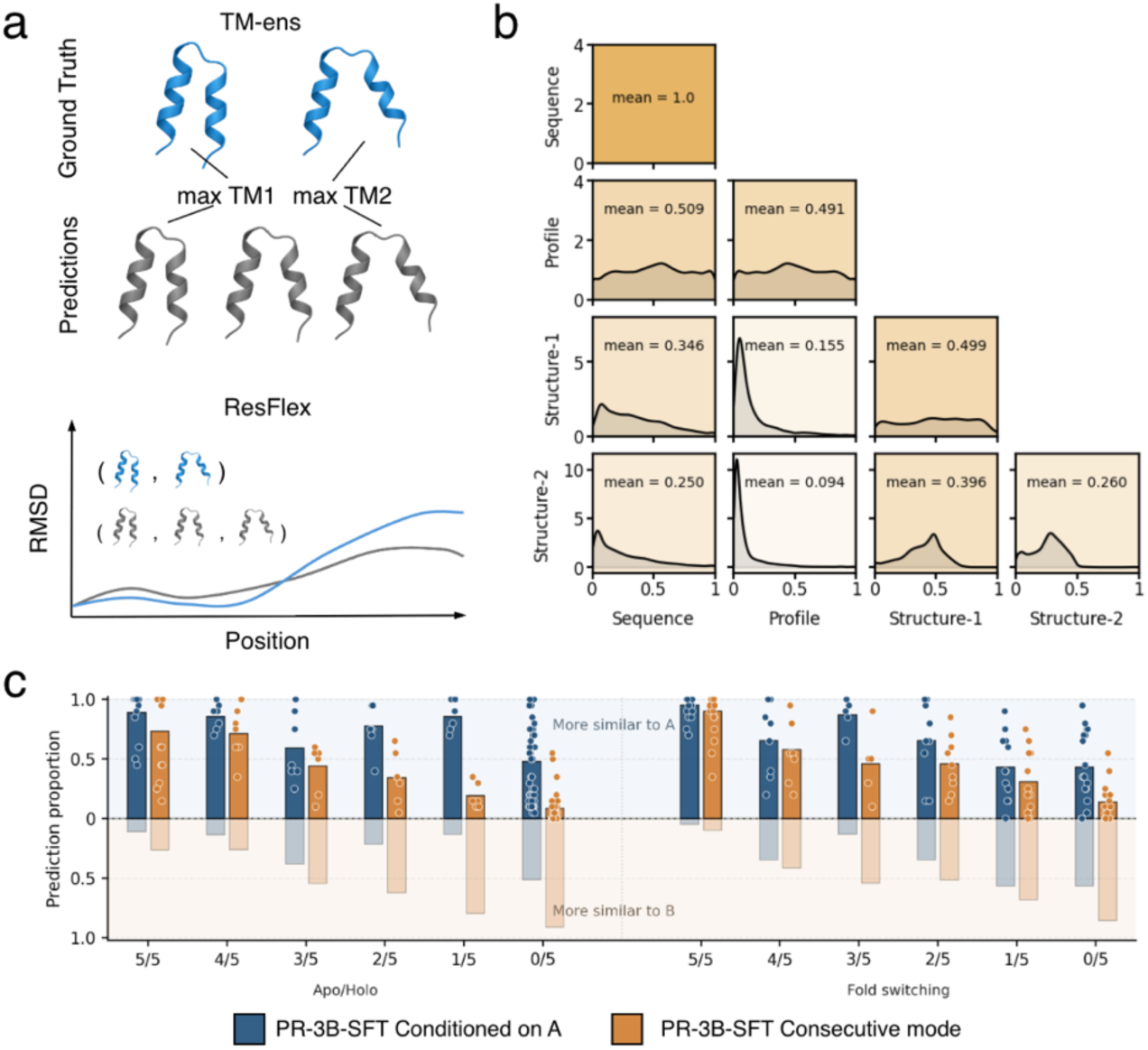
Analysis of modality-chain behavior in multiple-conformation prediction. **a**, Illustration of the evaluation metrics. Ensemble TM-score (TM-ens; top) measures structural accuracy through maximum TM-score matching between predicted and ground-truth conformations. Residue flexibility (ResFlex; bottom) measures agreement between their per-residue RMSD profiles. **b**, Cross-modality attention distributions in multiple-conformation prediction chain. Notable attention mass to the first structure is consistent with cross-conformation conditioning. **c**, Comparison of ProteinReasoner-3B-SFT conditioned on a given conformation A with consecutive generation. Proteins are grouped by similarity votes from five structures generated by pretrained ProteinReasoner, ranging from 5/5 predictions closer to conformation A to 0/5. Bars show the proportions of outputs closer to conformations A and B, and points denote individual proteins, for the apo/holo (left) and fold-switching (right) benchmarks. Conditioning on conformation A increased the proportion of outputs closer to A across all pretrained preference groups, supporting effective conditioning of the second conformation on the first.

**Extended Data Fig. 5.**
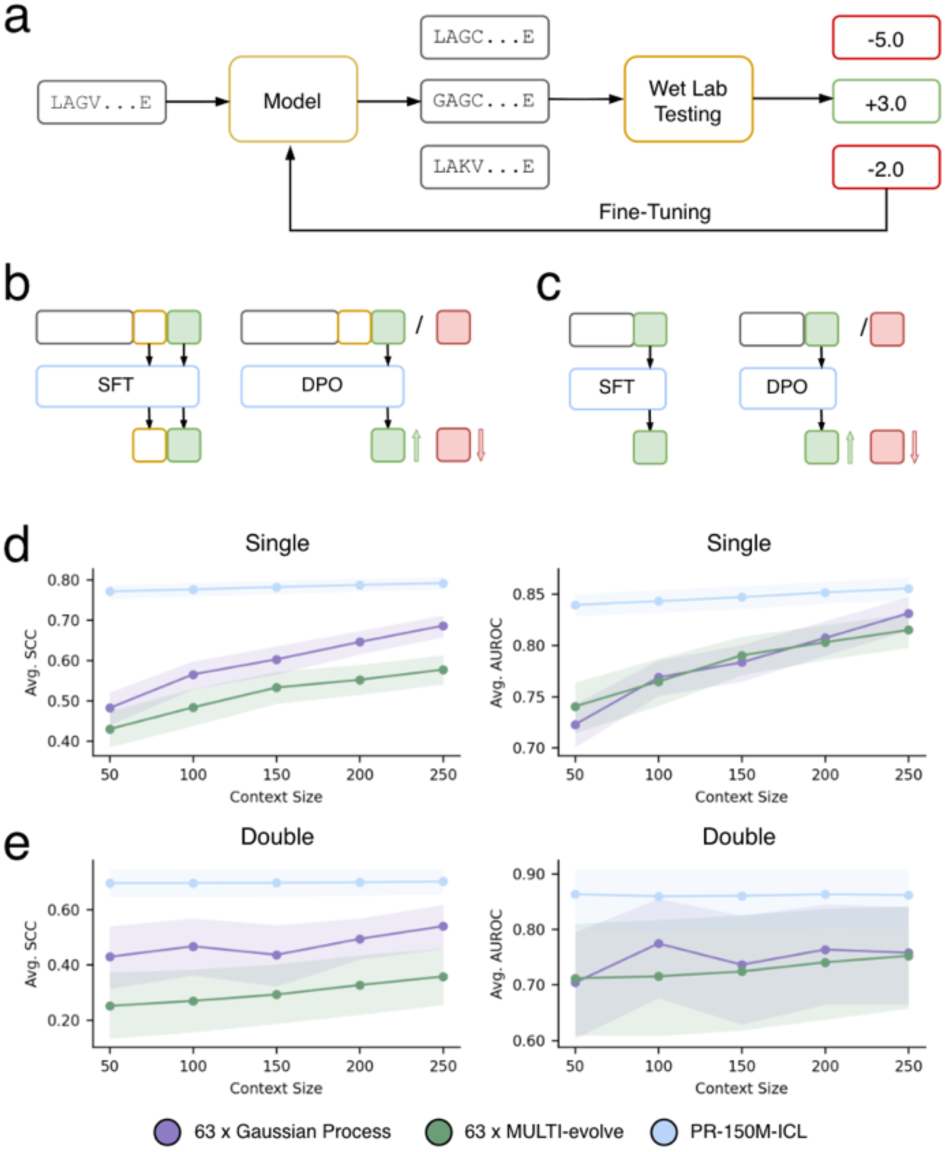
Training formulations and context-size analysis for protein optimization. **a**, Active learning (AL) workflow in which experimentally measured mutant sequences are used to update a target-specific model for subsequent rounds of candidate generation. **b**, Supervised fine-tuning (SFT) and direct preference optimization (DPO) of the ProteinReasoner in-context learning (ICL) model using the extended modality chain containing a directed evolution (DE) profile. SFT supervises reconstruction of the supplied DE profile and prediction of the corresponding mutant sequence, whereas DPO increases the relative score of preferred over less-preferred mutant sequences under the same context. **c**, SFT and DPO of the fine-tuned ProteinReasoner baseline, which retains the original structure–profile–sequence chain and does not use in-context experimental feedback. **d,e**, Average Spearman correlation coefficient (SCC) and area under the receiver operating characteristic curve (AUROC) as a function of context size for single mutants (**d**) and double mutants (**e**). PR-150M-ICL is compared with target-specific Gaussian process regression and MULTI-evolve baselines using 50– 250 experimentally scored context mutant sequences. Lines show mean performance across proteins and shaded regions indicate 95% bootstrap confidence intervals.

**Extended Data Fig. 6.**
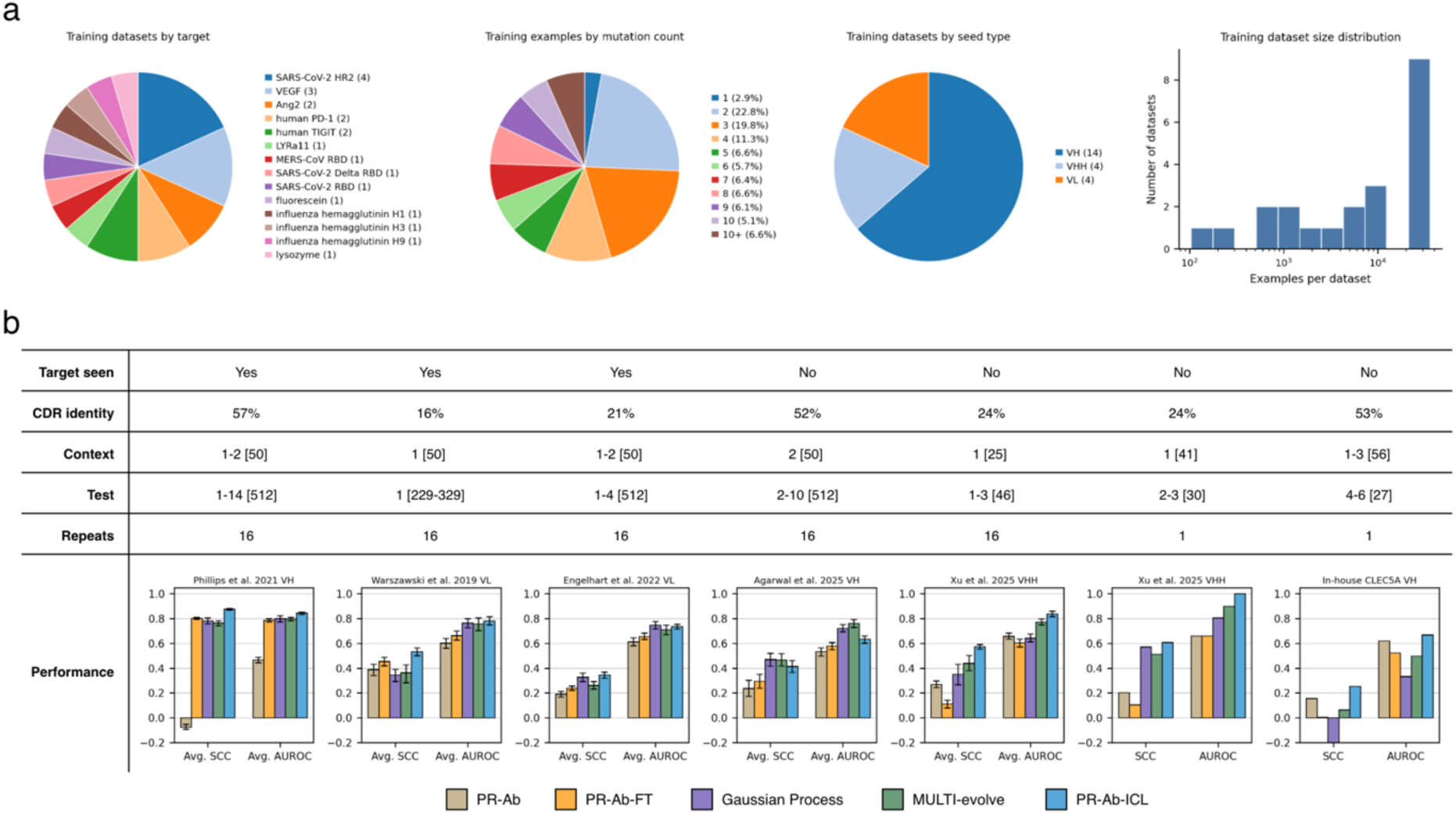
Affinity maturation datasets and benchmark design. **a**, Composition of the 22 affinity deep mutational scanning (DMS) training datasets by target, mutation depth, antibody or nanobody seed type, and dataset size. **b**, Summary of the seven affinity maturation test cases, including target representation in training, maximum complementarity-determining region (CDR) sequence identity to any training seed, context and test-set composition, number of context-sampling repeats, and performance of PR-Ab, PR-Ab-FT, Gaussian process regression, MULTI-evolve and PR-Ab-ICL. For cases with 16 context-sampling repeats, performance is reported as mean [95% bootstrap confidence interval] across repeats; cases evaluated once are reported as point estimates. Entries such as “1–2 [50]” denote 50 mutant sequences, each containing one to two mutations.

**Extended Data Fig. 7.**
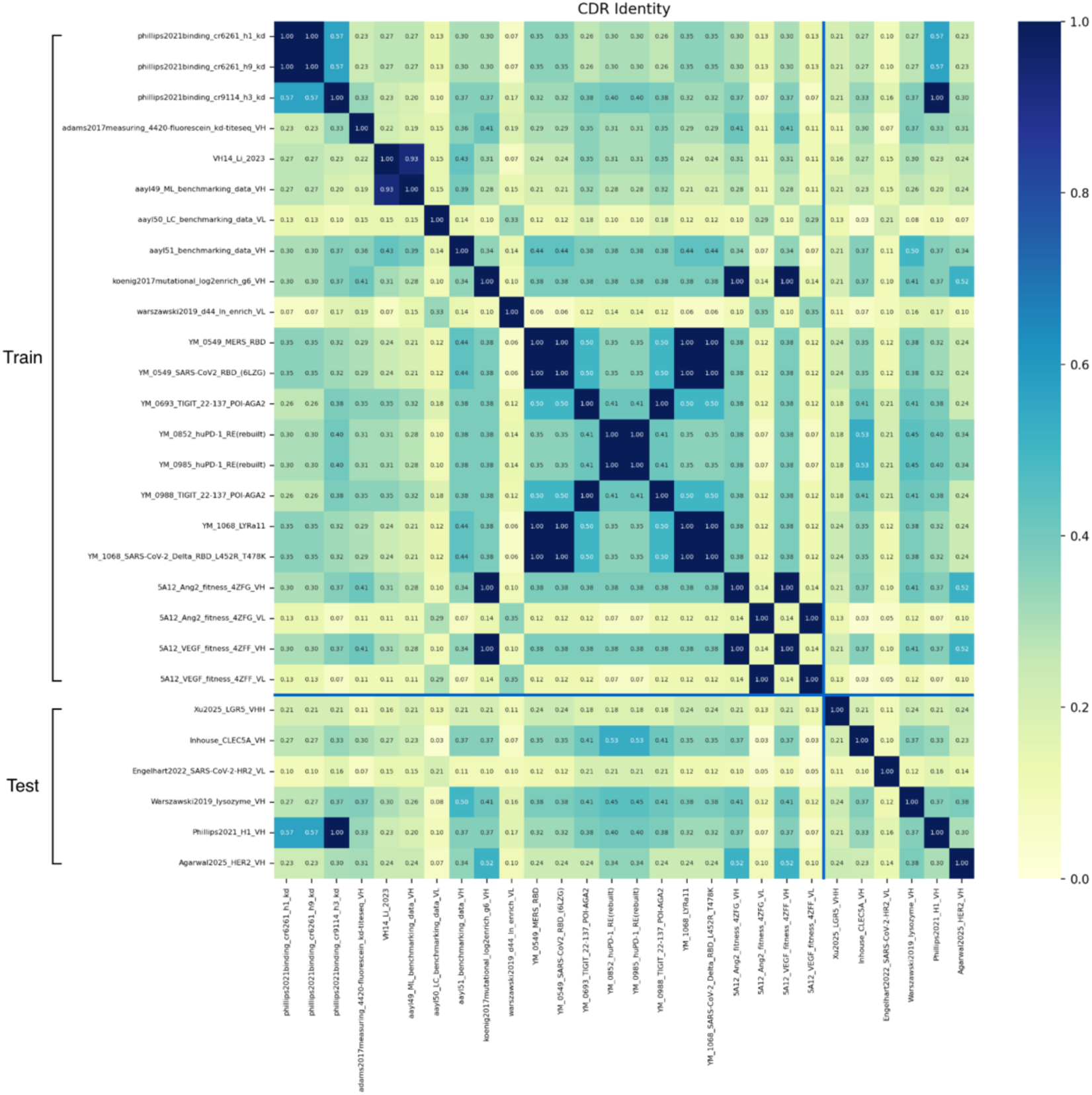
CDR sequence identity across affinity maturation datasets. Pairwise complementarity-determining region (CDR) sequence identity among the 22 training datasets and six held-out test seeds. Rows and columns are grouped by training or test status. Color denotes pairwise CDR sequence identity from 0 to 1.

