## Supplementary Information for "Modality-chain reasoning enables multimodal protein modelling and design"

#### Supplementary Results

##### Ablation studies of modality-chain design and pretraining strategies

We evaluated several ProteinReasoner-150M ablations to distinguish the contribution of the intermediate evolutionary profile from the effects of training direction and two-stage initialization. All models were trained on the same multimodal dataset and evaluated after 60,000 optimization steps; hyperparameters are summarized in **Supplementary Table 1**. The corresponding architectures and complete results are shown in **Extended Data Fig. 2**.

Including the evolutionary profile improved structure prediction and fitness prediction relative to the end-to-end model trained directly between sequence and structure. In structure prediction, ProteinReasoner-150M improved TM-score by 15.9–29.7% in the externally guided setting and by 4.8–8.2% in the internally inferred setting. Replacing the evolutionary profile with a uniform, non-informative matrix also reduced performance, indicating that the benefit arose from the evolutionary information represented by the profile rather than from simply inserting an additional modality segment (**Extended Data Fig. 2d,e**). On ProteinGym, ProteinReasoner-150M improved Spearman correlation by 26.7% relative to the end-to-end baseline score of 0.386 (**Extended Data Fig. 2h**). In inverse folding, the end-to-end model achieved comparable performance and marginally higher amino acid recovery on some datasets, whereas ProteinReasoner achieved similar or higher self-consistency TM-score. Thus, the profile-mediated chain did not uniformly maximize residue recovery but better preserved structural compatibility of the generated sequences (**Extended Data Fig. 2f,g**).

Training direction affected performance asymmetrically across tasks. The forward-only model, trained exclusively on sequence → evolutionary profile → structure, outperformed the bidirectionally trained ProteinReasoner on structure prediction, indicating a benefit from task-aligned specialization in this direction. However, inverse-only pretraining did not yield analogous gains in inverse folding: ProteinReasoner, the End-to-End model and the inverse-only model achieved comparable scTM, while the inverse-only model showed comparable or lower AAR across benchmarks. By contrast, bidirectional ProteinReasoner achieved the strongest ProteinGym performance. Together, these results indicate that restricting training to a single chain can improve structure prediction but does not universally benefit the corresponding reverse task, whereas exposure to both directions better supports mutation-effect prediction.

Two-stage initialization was also important. Across structure prediction and inverse folding, models initialized from the sequence-only pretrained backbone outperformed otherwise matched models trained from scratch. The difference was largest in the internally inferred setting, in which the model had to generate the evolutionary profile before producing the target modality, indicating that large-scale sequence pretraining supports effective internal profile generation. The benefit was smaller on ProteinGym but remained positive overall. Together, these ablations support both the use of an explicit evolutionary-profile intermediate and the two-stage pretraining strategy, while showing that the optimal directionality depends on the downstream task.

#### Supplementary Methods

##### Ablation models and training

We constructed a series of ProteinReasoner-150M ablations to evaluate the contribution of modality-chain composition, evolutionary-profile information and two-stage initialization. Unless otherwise specified, all ablation models were trained on the same multimodal pretraining dataset for 60,000 optimization steps and were evaluated at the same checkpoint. Hyperparameters not directly involved in an ablation were matched across models; learning rates were adjusted only where necessary to avoid early overfitting (**Supplementary Table 1**).

The forward-only model was trained only on the sequence → evolutionary profile → structure chain, whereas the inverse-only model was trained only on the structure → evolutionary profile → sequence chain. The from-scratch model used the same bidirectional modality chains as ProteinReasoner but did not inherit the sequence embedding, Transformer backbone or sequence output head from the sequence-only first-stage model. The end-to-end model removed the evolutionary-profile segment and was trained directly on sequence → structure and structure → sequence chains. For the uniform-profile evaluation, the evolutionary profile supplied to ProteinReasoner was replaced by a non-informative matrix in which all 21 residue states, including the gap state, had probability $1/21$ at every position. Ablation results are reported in **Extended Data Fig. 2**.

##### Detailed multimodal pretraining objective

For an autoregressive modality chain, ProteinReasoner predicts each target position from the preceding tokens under a causal mask. For the sequence and structure modalities, supervision is applied to all L residue tokens and the end-of-sequence (EOS) token, whereas the evolutionary-profile loss is applied only to the L residue-aligned profile positions and does not supervise an EOS token. Let $I_{\text{seq}}$, $I_{\text{struct}}$ denote the supervised sequence and structure positions, including the EOS position, and $I_{\text{profile}}$ denote the $L$ evolutionary-profile positions. The discrete sequence and structure cross-entropy losses were

$$\mathcal{L}_{\text{seq}} = -\frac{1}{L+1}\sum_{t\in I_{\text{seq}}} \log p_\theta\,(x_t \mid x_{<t}),$$

$$\mathcal{L}_{\text{struct}} = -\frac{1}{L+1}\sum_{t\in I_{\text{struct}}} \log p_\theta\,(x_t \mid x_{<t}).$$

For an evolutionary profile $P \in \mathbb{R}^{L\times 21}$ and its predicted distribution $\hat{P}$, the profile loss was

$$\mathcal{L}_{\text{prof}} = \frac{1}{L}\sum_{i=1}^{L} \text{KL}\,\left(P_i \,\|\, \hat{P}_i\right).$$

The total pretraining loss was

$$\mathcal{L} = \lambda_{\text{seq}}\mathcal{L}_{\text{seq}} + \lambda_{\text{struct}}\mathcal{L}_{\text{struct}} + \lambda_{\text{prof}}\mathcal{L}_{\text{prof}},$$

with model-specific modality weights reported in **Supplementary Table 3**.

Large-scale training used a customized Megatron–DeepSpeed implementation with data parallelism across

GPUs. Optimization and hardware settings are described in the main Methods, and model-specific learning rates, warm-up proportions and modality-loss weights are reported in **Supplementary Table 3**.

##### Multiple-conformation fine-tuning objective

For multiple-conformation prediction, the input chain was sequence → evolutionary profile → structure$_1$ → structure$_2$. Supervision was applied to both structure-token segments. If $I_{s_1}$ and $I_{s_2}$ denote the residue-token positions of structure$_1$ and structure$_2$, respectively, the fine-tuning objective was

$$\mathcal{L}_{\mathrm{multi-conf}} = -\frac{1}{L}\sum_{t\in I_{s_1}} \log p_\theta\left(x_t \mid x_{<t}\right) - \frac{1}{L}\sum_{t\in I_{s_2}} \log p_\theta\left(x_t \mid x_{<t}\right).$$

Training configurations for the two-structure chain and the corresponding one-structure control models are reported in **Supplementary Table 4**.

##### Directed evolution profile construction

Given $K$ experimentally characterized context sequences $\left\{\left(s^{(k)}, y_k\right)\right\}_{k=1}^{K}$, where each sequence has length $L$ and each experimental property value is centred on the corresponding wild type and z-score normalized, we constructed a directed evolution (DE) profile over the 20 standard amino acids.

Each context sequence was represented by a one-hot feature map

$$\phi\left(s^{(k)}\right) \in \{0,1\}^{L\times 20},$$

$$\phi\left(s^{(k)}\right)_{i,a} = \mathbf{1}\left(s_i^{(k)} = a\right).$$

The Hamming-similarity Gram matrix $K_H \in \mathbb{R}^{K\times K}$ was defined as

$$(K_H)_{kk'} = \left\langle \phi\left(s^{(k)}\right), \phi\left(s^{(k')}\right)\right\rangle = \sum_{i=1}^{L} \mathbf{1}\left(s_i^{(k)} = s_i^{(k')}\right) = L - d_H\left(s^{(k)}, s^{(k')}\right),$$

where $d_H$ denotes Hamming distance. Ordinary-kriging weights $w \in \mathbb{R}^K$ were obtained from

$$\begin{bmatrix} K_H + \lambda I & \mathbf{1} \\ \mathbf{1}^\top & 0 \end{bmatrix} \begin{bmatrix} w \\ \mu \end{bmatrix} = \begin{bmatrix} y \\ 1 \end{bmatrix}$$

where $y = (y_1, \ldots, y_K)^\top$, $\mu$ is the Lagrange multiplier enforcing $\sum_k w_k = 1$, and the ridge coefficient was set to

$$\lambda = 0.1 \ \mathrm{Var}\left(\{y_{\mathrm{WT}}\} \cup \{y_k\}_{k=1}^{K}\right),$$

calculated after wild-type centering and z-score normalization.

The fitted sequence weights were projected back into residue space to form an unnormalized profile $U \in \mathbb{R}^{L\times 20}$:

$$U_{i,a} = \sum_{k=1}^{K} w_k \, \mathbf{1}\left(s_i^{(k)} = a\right).$$

Because kriging coefficients can be negative, a position-wise softmax was applied to obtain the DE profile used by ProteinReasoner:

$$D_{i,a} = \frac{\exp\left(U_{i,a}\right)}{\sum_{a'=1}^{20} \exp\left(U_{i,a'}\right)}.$$

Thus, $D_{i,a} \geq 0$ and $\sum_a D_{i,a} = 1$ at every position. Profile construction was performed offline during dataset preparation, and the resulting $L \times 20$ tensor was provided directly to ProteinReasoner during training and inference.

Before softmax normalization, the additive score of a query sequence $q$ can be written as

$$\tilde{f}(q; U) = \sum_{i=1}^{L} U_{i,q_i} = k_H(q, S)^{\mathsf{T}} w,$$

where $k_H(q, S)$ contains the Hamming similarities between $q$ and the context sequences. This is the ordinary-kriging or Gaussian-process posterior predictor under the Hamming-similarity kernel, up to the ordinary-kriging mean constraint. The DE profile therefore projects a kernel predictor into a residue-wise representation that can be processed using the same interface as a natural evolutionary profile.

**Affinity-specific conditioning modules**

The affinity-maturation models included a lightweight profile-conditioned causal cross-attention module between the Transformer backbone and the mutant-sequence output head. This module was not used in the Megascale thermostability experiments. Cross-attention was applied only to the $L$ residue positions, excluding modality-boundary tokens. DE-profile hidden states served as queries, and mutant-sequence hidden states served as keys and values. A causal mask restricted DE-profile position $i$ to mutant-sequence positions up to $i$. The cross-attention output was added residually to the corresponding mutant-sequence hidden states and followed by layer normalization; boundary-token representations were left unchanged. The module comprised two cross-attention layers, each with 16 attention heads and dropout of 0.1. Attention output projections were initialized to zero so that the module introduced minimal perturbation at the beginning of fine-tuning.

During DPO, the affinity-maturation ICL model additionally fused the mutant-sequence distribution with the input DE profile through a case-level gated correction in amino-acid log-probability space. Let $z_{i,a}$ denote the model logit and $P_{\mathrm{DE}}(i, a)$ the DE-profile probability for amino acid $a$ at position $i$. The fused distribution over the 20 canonical amino acids was

$$P_{\mathrm{fused}}(i, a) = \mathrm{softmax}_{a \in \mathcal{A}}\left[z_{i,a} + \lambda(C) \log P_{\mathrm{DE}}\,(i, a)\right].$$

The fusion coefficient was shared across residue positions and conditioned on the experimental context. DE-profile hidden states were mean-pooled over residue positions, excluding boundary tokens, to obtain $h_C$ and the coefficient was predicted as

$$\lambda(C) = \lambda_{\max}\sigma[W_2\,\mathrm{GELU}(W_1\,\mathrm{LN}(h_C)) + b_2],$$

with $\lambda_{\max} = 3$. The final projection weights were initialized to zero, and its bias was initialized such that $\lambda(C) = 1$ at the start of DPO training. Because $\lambda(C)$ depends only on the shared context, both sequences in each preference pair received the same gate value.

Fusion was applied only within the conditional distribution over canonical amino acids; probability mass assigned to amino acids relative to special tokens was preserved, and special-token probabilities were unchanged. EOS was excluded from sequence scoring. Policy and reference models were initialized from the same SFT checkpoint and used the same fusion architecture; the reference model was frozen, whereas the policy model and fusion head were optimized with separate learning rates.

**Protein optimization training-example construction**

Training examples were preconstructed for both conventional fine-tuning (FT) and in-context learning (ICL). FT examples used the structure → evolutionary profile → mutant sequence chain, whereas ICL examples used structure → evolutionary profile → wild-type sequence → DE profile → mutant sequence. In both settings, structure tokens and natural evolutionary profiles were derived from the corresponding wild-type protein.

For supervised fine-tuning, mutant sequences with experimentally measured scores above the corresponding wild type were used as targets. For DPO, preferred and less-preferred mutant sequences were drawn from the same wild-type protein and were required to differ by at least a predefined score threshold. For ICL examples, each sampled context set was converted into a DE profile and paired with a fixed number of SFT targets or DPO pairs. Dataset-construction hyperparameters are reported in **Supplementary Table 7**.

The Megascale dataset used the standard construction procedure, whereas the affinity-maturation datasets used an extended procedure for multi-point contexts, mutation-position coverage, mixing of primary and context targets, and score-gap-aware preference-pair sampling. Detailed procedures for constructing the protein-optimization training datasets are provided in **Algorithms 1** and **2** at the end of the Supplementary Information.

**Additional affinity-dataset processing**

Experimental scores were directionally harmonized so that larger values represented stronger binding. For the Warszawski et al. dataset distributed through AbBiBench, the processed scores were multiplied by -1 because their direction was reversed relative to the original study.

For the four high-throughput evaluation cases, context sets contained 50 single- and double-mutant sequences and were sampled to contain approximately 25% experimentally improved sequences where available. When fewer improved sequences were available, the remaining context positions were filled with lower-scoring sequences.

**Target-specific baseline implementations**

**ThermoMPNN and ThermoMPNN-D.** ThermoMPNN and ThermoMPNN-D were evaluated using their official repositories and checkpoints. Before benchmark evaluation, the implementations were checked by reproducing the prediction scores supplied with the official example data. Both models were then evaluated on the same Megascale test split used for ProteinReasoner.

**Gaussian-process baselines**. For each held-out Megascale protein, the same experimentally characterized context mutant sequences supplied to the ICL model were used to fit each Gaussian-process baseline, and the remaining mutant sequences were used for evaluation. Residue-level representations from the corresponding fine-tuned ProteinReasoner model were mean-pooled across sequence positions to obtain fixed-dimensional mutant-sequence embeddings. An exact Gaussian process with a constant mean function, linear covariance kernel and Gaussian observation likelihood was fitted independently for each protein by maximizing the exact marginal log-likelihood. Optimization used Adam with a learning rate of 0.1 for up to 200 iterations and early stopping with patience 20 and tolerance $10^{-4}$. Candidate mutant sequences were ranked by posterior predictive mean.

**MULTI-evolve**. MULTI-evolve was implemented using its official repository. Two input representations were evaluated: (1) a flattened position-wise one-hot representation of the full mutant sequence and (2) mean-pooled residue embeddings from ESM-2-3B (*esm2_t36_3B_UR50D*). For each target protein and representation, fully connected networks were evaluated over

- number of hidden layers: $\{1, 2, 3\}$;
- learning rate: $\{10^{-4}, 10^{-3}, 10^{-2}\}$;
- mini-batch size: $\{4, 8\}$.

This produced 18 configurations for each representation. All networks used 100 hidden units per layer, LeakyReLU activations with negative slope 0.2, dropout of 0.2 and the Adam optimizer. Models were trained for up to 300 epochs with early stopping after 15 epochs without validation improvement. Each configuration was trained under three-fold cross-validation, yielding three fold-specific models. The input representation and hyperparameter configuration with the lowest mean validation mean-squared error were selected independently for each target protein. Final predictions for the remaining mutant sequences were obtained by averaging predictions from the three corresponding fold-specific models.

##### Additional attention-analysis details

Tokens were assigned to modalities from their positions in the ordered input chain. Although the DE profile is a continuous rather than discrete modality, its $L \times D$ hidden representation was treated as a separate modality for attention analysis. Attention matrices were averaged across all Transformer layers and heads. For each sample, attention from a source modality to a target modality was summed over the corresponding query–key pairs and normalized by the total outgoing attention mass from the source modality. Modality-boundary tokens were excluded. Sample-level values were summarized across benchmark datasets to obtain the reported distributions and means.

#### Supplementary Tables

| Model | Loss Wts | Max LR | Min LR | Warmup | Steps | Seq Len |
|---|---|---|---|---|---|---|
| ProteinReasoner | 1,2,1 | 2e-3 | 2e-5 | 0.62% | 60k | 3072 |
| Forward | 1,2,1 | 2e-3 | 2e-5 | 0.62% | 60k | 3072 |
| Inverse | 1,2,1 | 2e-3 | 2e-5 | 0.62% | 60k | 3072 |
| FromScratch | 1,2,1 | 4e-4 | 1e-5 | 0.62% | 60k | 3072 |
| End-to-End | 1,1 | 2e-3 | 2e-5 | 0.62% | 60k | 2048 |

**Supplementary Table 1.** Ablation model hyperparameters. *Loss Wts:* Weights for structure, profile, and sequence losses; *Warmup:* Percentage of steps used for learning rate warmup; *Steps:* Total number of training steps.

| Model | Layers | Dim | FFN Dim | Heads | Dropout |
|---|---|---|---|---|---|
| ProteinReasoner-150M | 30 | 640 | 1706 | 20 | 0.1 |
| ProteinReasoner-650M | 33 | 1280 | 3424 | 20 | 0.1 |
| ProteinReasoner-3B | 36 | 2560 | 6832 | 40 | 0.1 |

**Supplementary Table 2.** Model architecture details. *Layers*: number of transformer layers; *Dim*: Hidden embedding dimension; *FFN Dim*: Feedforward layer intermediate dimension; *Heads*: number of attention heads.

| Model | Loss Wts | Max LR | Min LR | Warmup | Seq Len |
|---|---|---|---|---|---|
| ProteinReasoner-150M | 1,2,1 | 2e-3 | 2e-5 | 0.62% | 3072 |
| ProteinReasoner-650M | 1,1,1 | 2e-3 | 2e-5 | 1.25% | 3072 |
| ProteinReasoner-3B | 1,2,1 | 5e-4 | 1e-5 | 1.25% | 3072 |

**Supplementary Table 3.** Pretraining hyperparameters. *Loss Wts:* Weights for structure, profile, and sequence losses; *Warmup:* Percentage of steps used for learning rate warmup.

| | Chain | Size | LR | Epochs | Batch Size |
|---|---|---|---|---|---|
| 1 | Seq-Prof-Struct-Struct | 650M | 1e-5 | 1 | 128 |
| 2 | Seq-Prof-Struct-Struct | 3B | 2e-5 | 2 | 128 |
| 3 | Seq-Prof-Struct | 650M | 1e-5 | 1 | 128 |
| 4 | Seq-Prof-Struct | 3B | 1e-5 | 2 | 128 |

**Supplementary Table 4.** Multiple-conformation prediction fine-tuning hyperparameters. *Chain*: modality-chain configuration.

| Model | Task | Optimizer | LR | LR Schedule | Warmup | Min LR | Batch size | Steps |
|---|---|---|---|---|---|---|---|---|
| PR-650M | Comparison to ThermoMPNN | AdamW | 1e-5 | Cosine | 2% | 1e-6 | 64 | 1190 |
| PR-150M-FT | Thermostability | AdamW | 1e-5 | Cosine | 2% | 1e-6 | 64 | 160 |
| PR-150M-ICL | Thermostability | AdamW | 1e-5 | - | - | 1e-5 | 64 | 8216 |
| PR-Ab-FT | Affinity | AdamW | 1e-5 | Cosine | 10% | 5e-6 | 128 | 400 |
| PR-Ab-ICL | Affinity | AdamW | 1e-5 | Cosine | 10% | 5e-6 | 128 | 400 |

**Supplementary Table 5.** Supervised fine-tuning configurations for ProteinReasoner protein optimization models.

| Model | Task | Optimizer | beta | LR | LR Schedule | Warmup | Min LR | Batch size | Steps |
|---|---|---|---|---|---|---|---|---|---|
| PR-650M | Comparison to ThermoMPNN | AdamW | 0.05 | 1e-5 | Cosine | 3% | 1e-6 | 64 | 9900 |
| PR-150M-FT | Thermostability | AdamW | 0.1 | 1e-5 | - | - | 1e-5 | 64 | 8220 |
| PR-150M-ICL | Thermostability | AdamW | 0.1 | 1e-5 | - | - | 1e-5 | 64 | 8220 |
| PR-Ab-FT | Affinity | AdamW | 0.1 | 1e-5 | Cosine | 1% | 2e-6 | 128 | 1408 |
| PR-Ab-ICL | Affinity | AdamW | 0.1 | 1e-5 | Cosine | 1% | 2e-6 | 128 | 1408 |

**Supplementary Table 6.** Directed preference optimization configurations for ProteinReasoner protein optimization models.

| Dataset | N | R | T (SFT / DPO) | g | ρ | Gap bins | Bin weights |
|---|---|---|---|---|---|---|---|
| Thermostability PR-FT in comparison with ThermoMPNN | - | - | 320/256 | 0.5 | 0 | - | - |
| Thermostability PR-FT as baseline model for ICL | - | - | 640/1280 | 0.001 | 0 | - | - |
| Thermostability PR-ICL | 50 | 64 | 10/10 | 0.001 | 0 | - | - |
| Affinity maturation dataset (FT and ICL share the same) | 50 | 64 | 128/128 | 0.05 | 0.1 | [0.05, 0.1, 0.2, 1.0, 100.0] | [1.0, 2.0, 2.0, 1.0] |

**Supplementary Table 7.** Dataset-construction hyperparameters for ProteinReasoner protein optimization models. $N$: context size; $R$: number of contexts redraws per seed; $T$: number of targets/pair per context or seed; $g$: minimum preference pair gap; $\rho$: ratio of context sequence as target.

### Dataset Construction Algorithms

The two algorithms below describe the in-context dataset builders used for training. For each wild-type (WT) case, a set of experimentally-scored single/multi-point mutants is sampled to form an *in-context demonstration set*, which is summarized into a length-$L$ *directed-evolution (DE) profile* $P \in \mathbb{R}^{L\times 20}$ via Gaussian-process (Kriging) regression over a full-length Hamming similarity kernel. The model is then asked to generate *target* mutant sequences (SFT) or to rank *preference pairs* of mutants (DPO), conditioned on the same DE profile.

**Notation.** $D$ is the set of WT (seed) proteins in a split. $M = \{(m, s)\}$ is the pool of measured mutants for a WT case, where $m$ is a mutation string (e.g. `A12G` or `A12G,K45R`) and $s$ its assay score; $s_{\text{wt}}$ is the WT score. BuildDEProfile$(\cdot)$ returns the Kriging profile from a set of scored context sequences (ordinary Kriging, full-length Hamming kernel). mut_count$(m)$ is the number of point mutations in $m$ and pos$(m)$ their positions. Tokenize$(\cdot)$ is the sequence tokenizer producing model input tokens.

**Algorithm 1** Megascale training dataset builder (SFT and DPO).
Hyper-parameters: context size $N = 50$, redraws of context $R = 64$, targets-per-context $T = 10$, pair gap $g = 0.001$.

**Input:** seed proteins $D$; for each, sequence $x$, structure tokens *Struct*, Evolutionary Profile $P_{evo}$ mutant pool $M = \{(m, s)\}$, WT score $s_{\text{wt}} = 0$
**Params:** context size $N$, redraws $R$, targets-per-context $T$, pair gap $g$ (DPO)
**Output:** list of training examples

**for** each seed protein in $D$ **do**
  $M_{\text{single}} \leftarrow \{(m, s) \in M : \text{mut\_count}(m) = 1\}$ ▷ *single mutants*
  **for** rep $= 1$ **to** $R$ **do**
    *// In-context demonstration set + DE profile*
    $C \leftarrow$ RandomSample$(M_{\text{single}}, \min(N, |M_{\text{single}}|))$
    $P \leftarrow$ BuildDEProfile$(C \cup \{\text{WT}\})$ ▷ *Kriging, $\mathbb{R}^{L\times 20}$*
    $M_{\text{rest}} \leftarrow \{(m, s) \in M : m \notin C\}$ ▷ *not used as context*
    **if** SFT **then**
      pool $\leftarrow \{(m, s) \in M_{\text{rest}} : s > s_{wt}\}$ ▷ *stabilizing*
      targets $\leftarrow$ RandomSample(pool, $\min(T, |\text{pool}|)$)
      **for** $(m, s) \in$ targets **do**
        **emit** $\{$*Struct*, $P_{evo}$, $x$, $P$, Tokenize(ApplyMutation$(x, m)$), $s\}$
    **else if** DPO **then**
      $M_{\text{pair}} \leftarrow M_{\text{rest}} \cup \{(\text{WT}, 0)\}$
      **for** $k = 1$ **to** $T$ **do**
        **repeat** (up to `max_retry`)
          $(a, b) \leftarrow$ RandomSample$(M_{\text{pair}}, 2)$
        **until** $|s_a - s_b| \geq g$ ▷ *require score gap*
        $(w, l) \leftarrow (a, b)$ if $s_a > s_b$ else $(b, a)$ ▷ *winner/loser*
        **emit** $\{$*Struct*, $P_{evo}$, $x$, $P$, Tokenize$(w)$, Tokenize$(l)$, $s_a$, $s_b\}$
**return** examples

The DE profile $P$ is computed once per redraw and reused across all $T$ targets/pairs, so the model sees each context paired with multiple outputs at amortized cost. Evaluation reuses the same builder with score type `logits_deprofile` (context ordered by a pretrained model's log-likelihood) and one deterministic target draw per case.

**Algorithm 2** Affinity-maturation training dataset builder (SFT and DPO).
Extensions over Alg. 1: (i) multi-point context (1–3 mutations); (ii) coverage-restricted targets; (iii) primary plus context targets mixed by ratio $\rho$; (iv) gap-binned, reuse-capped preference pairing.

**Input:** seed proteins $D$; for each, sequence $x$, structure tokens $\mathit{Struct}$, Evolutionary Profile $P_{evo}$, mutant pool $M = \{(m, s)\}$, WT score $s_{\text{wt}}$
**Params:** context size $N$, min context $N_{\min} = 24$, mutation range $[c_{\text{lo}}, c_{\text{hi}}] = [1, 3]$, redraws of context $R = 64$, target ratio $\rho = 0.1$, max targets $T_{\max} = 128$; *(DPO)* gap bins $B$, bin weights $W$, min gap $g_{\min} = 0.05$, context-pair ratio $\rho_p = 0.1$, reuse cap $\kappa = 8$, max pairs $E = 128$
**Output:** list of training examples

**for** each seed protein in $D$ **do**
  **for** rep $= 1$ **to** $R$ **do**
    *// (1) Select context demonstration set*
    $E_{\text{pool}} \leftarrow \{(m, s) \in M : c_{\text{lo}} \leq \text{mut\_count}(m) \leq c_{\text{hi}}\}$
    **if** $|E_{\text{pool}}| < N_{\min}$ **then continue**
    $C \leftarrow$ RandomSample($E_{\text{pool}}$, $\min(N, |E_{\text{pool}}|)$)
    $C \leftarrow C \cup \{(\text{WT}, s_{\text{wt}})\}$ ▷ *append WT*
    *// (2) Build DE profile (Kriging)*
    $P \leftarrow$ BuildDEProfile($C$); cov $\leftarrow \bigcup_{(m,\cdot) \in C} \text{pos}(m)$
    *// (3) Build target pools (targets must be coverage-restricted)*
    Cand $\leftarrow \{(m, s) \in M : m \notin C \text{ and } \text{pos}(m) \subseteq \text{cov}\}$
    $T_{\text{ctx}} \leftarrow$ {context mutants with $s > s_{\text{wt}}$} ▷ *context as target*
    **if** SFT **then**
      $T_{\text{pri}} \leftarrow \{(m, s) \in \text{Cand} : s > s_{\text{wt}}\}$ sorted desc. ▷ *higher than WT*
      *// (4a) mix primary + context targets by ratio $\rho$*
      targets $\leftarrow$ LimitTargetsPerContext($T_{\text{pri}}, T_{\text{ctx}}, T_{\max}, \rho$)
      **for** $t \in$ targets **do**
        **emit** {$\mathit{Struct}$, $P_{evo}$, $x$, $P$, Tokenize($t$), $s_t$}
    **else if** DPO **then**
      $T_{\text{pri}} \leftarrow$ Cand sorted desc. ▷ `all`
      *// (4b) gap-binned, reuse-capped layered pairing*
      $\text{Cand}_{\text{pairs}} \leftarrow$ Unique($T_{\text{pri}} \cup T_{\text{ctx}}$)
      **if** $|\text{Cand}_{\text{pairs}}| < 2$ **then continue**
      Bins $\leftarrow$ BinPairs($\text{Cand}_{\text{pairs}}, B, g_{\min}$) ▷ *ordered $(w, l)$, $s_w > s_l$, drop gap $\leq g_{\min}$*
      pairs $\leftarrow \emptyset$
      **while** $|\text{pairs}| < E$ **and** any bin non-empty **do**
        group $\leftarrow$ WeightedChoice(context, primary) ▷ *by remaining $\rho_p$ quotas*
        $b \leftarrow$ WeightedChoice(non-empty bins of group; $W$)
        $(w, l, \text{gap}) \leftarrow$ PopPair(Bins[group][$b$])
        **if** reuse($w$) $\geq \kappa$ **or** reuse($l$) $\geq \kappa$ **then continue**
        pairs $\leftarrow$ pairs $\cup \{(w, l, \text{gap})\}$; increment reuse($w$), reuse($l$)
      **for** $(w, l, \text{gap}) \in$ pairs **do**
        **emit** {$\mathit{Struct}$, $P_{evo}$, $x$, $P$, Tokenize($w$), Tokenize($l$), gap}
**return** examples
